# The gut hormone Allatostatin C regulates food intake and metabolic homeostasis under nutrient stress

**DOI:** 10.1101/2020.12.05.412874

**Authors:** Olga Kubrak, Line Jensen, Nadja Ahrentløv, Takashi Koyama, Alina Malita, Muhammad T. Naseem, Mette Lassen, Stanislav Nagy, Michael J. Texada, Kenneth A. Halberg, Kim Rewitz

## Abstract

The intestine is a central regulator of metabolic homeostasis. Dietary inputs are absorbed through the gut, which senses their nutritional value and relays hormonal information to other organs to coordinate systemic energy balance. However, the specific gut hormones that communicate energy availability to target organs to induce appropriate metabolic and behavioral responses are poorly defined. Here we show that the enteroendocrine cells (EECs) of the *Drosophila* gut sense nutrient stress via the intracellular TOR pathway, and in response secrete the peptide hormone allatostatin C (AstC). Gut-derived AstC induces secretion of glucagon-like adipokinetic hormone (AKH) via its receptor AstC-R2, a homolog of mammalian somatostatin receptors, to coordinate food intake and energy mobilization. Loss of gut *AstC* or its receptor in the AKH-producing cells impairs lipid and sugar mobilization during fasting, leading to hypoglycemia. Our findings illustrate a nutrient-responsive endocrine mechanism that maintains energy homeostasis under nutrient-stress conditions, a function that is essential to health and whose failure can lead to metabolic disorders.

## Introduction

Organisms’ survival depends on their ability to maintain energetic homeostasis^1^, and deficiency in this ability is implicated in the pathogenesis of human diseases such as diabetes and obesity. Maintaining whole-body energy balance requires coordination of nutrient sensing, energy metabolism, and nutrient intake, functions distributed across discrete organs. Such coordination is achieved through the exchange of metabolic information between organs, mediated by nutrient-responsive hormones. These hormones are secreted by specialized tissues that sense changes in internal and external nutritional conditions, and underlie inter-tissue communication that maintains systemic energy homeostasis via the regulation of appetite and metabolism. Defects in this exchange of nutritional information between tissues can lead to metabolic dysfunction. Thus, characterizing the network of hormones that convey nutrient availability and govern energy homeostasis through the control of food intake and energy expenditure is critical for understanding human metabolic disorders and for developing future therapeutics.

The gut is the largest endocrine organ and a central coordinator of systemic energy homeostasis^2^. Ingested nutrients are absorbed through this organ, which also senses nutritional value and relays metabolic information to other tissues via secretion of gut hormones that are released into circulation from enteroendocrine cells (EECs) and coordinate body-wide processes^3^. The EECs therefore play an important role in balancing energy homeostasis through crosstalk with regulatory networks that regulate appetite and metabolism. In the mammalian intestine, for example, the EECs release ghrelin during fasting, which signals directly to the brain to regulate appetite and energy homeostasis^4^. Glucagon-like peptide 1 (GLP-1) is secreted by the EECs in a glucose-dependent manner and plays a key role in regulating glucose metabolism^5^. Together GLP-1 and gastric inhibitory peptide (GIP) underlie the so-called incretin effect, through which oral glucose consumption induces a much stronger insulin-secretory response than intravenous administration of glucose; GLP-1 is secreted from the intestine in response to ingestion of glucose and acts directly on the pancreatic *β*-cells to potentiate glucose-dependent insulin secretion. This incretin effect is diminished in type-2 diabetes patients, and the use of incretin-hormone therapy is of great medical important in the treatment of metabolic disorders.

The *Drosophila* gut is emerging as a central paradigm for studies of the systemic effects of intestinal homeostasis. The fly gut shares structural and functional features with the mammalian intestine, and many of the systems that mediate nutrient signaling and control metabolism are evolutionarily ancient and conserved^1,6,7^. In *Drosophila*, metabolic homeostasis is regulated by hormones similar to insulin and glucagon. In mammals, insulin promotes the cellular uptake and conversion of excess nutrients into glycogen and lipids, which are stored in liver and adipose tissues under nutrient-replete conditions. Under conditions of nutrient scarcity, mammalian glucagon induces the mobilization of stored energy to provide the means to sustain glycemic control and basic physiological body functions. Likewise, flies express several conserved insulin-like peptides (DILPs) in insulin-producing cells (IPCs) of the brain, which are believed to be functionally and evolutionarily related to mammalian pancreatic *β* cells^8,9^. During nutrient abundance, these DILPs promote growth and metabolism through their activation of a single insulin receptor. Adipokinetic hormone (AKH) is functionally related to mammalian glucagon and is known to act antagonistically to insulin. During nutritional scarcity, AKH is secreted from AKH-producing cells (APCs) of the corpora cardiaca (CC), neurosecretory tissues analogous with pancreatic *α*-cells, to promote energy mobilization and food-seeking behaviors. The *Drosophila* fat body is the main energy storage organ and is functionally analogous to the mammalian liver and adipose tissues. During nutrient-stress conditions, AKH acts on the fat body via its receptor AkhR to promote the breakdown of stored lipids and carbohydrates to prevent hypoglycemia^10^. AKH also acts on a small set of arousal-promoting neurons in the brain to promote food-seeking behavior^11^.

Like the mammalian intestine, the EECs of the *Drosophila* gut secrete a cocktail of endocrine factors into the hemolymph (insect circulatory fluid) that relay information to other target organs. Although evidence suggests that both mammalian and *Drosophila* guts produce a large number of hormones, the functions of these signals, as well as the information they convey, remain poorly defined. In the adult fly gut, EECs are marked by expression of the transcription factor Prospero (Pros) and produce more than a dozen known peptide hormones^12-15^. These include Allatostatin A (AstA), AstC, Bursicon alpha (Burs), neuropeptide F (NPF), short neuropeptide F (sNPF), diuretic hormone 31 (Dh31), pigment-dispersing factor (PDF), CCHamides 1 and 2, Myoinhibiting peptide precursor (MIP or AstB), Orcokinin B, and Tachykinin (TK). However, the function of these gut hormones is known in only a few cases. Gut-derived TK regulates intestinal lipid homeostasis^16^, and NPF from the gut governs mating-induced germline stem-cell proliferation^17^. Burs is released from EECs in response to nutrient intake and regulates energy homeostasis through a neuronal relay that modulates AKH signaling^18^. Dh31 upregulates gut motility in response to enteropathic infection^19^.

Although incretin hormones and insulin secretion have been extensively studied in both mammals and *Drosophila*, the regulatory mechanisms controlling the secretion of glucagon remain poorly defined, and nutrient-responsive gut hormones with the ability to promote glucagon secretion are not known. To characterize factors secreted by the intestine that are important for energy homeostasis, we performed a genetic screen for gut hormones that affect metabolic stress responses. We show here that intestinal nutrient signaling is mediated by AstC and AstC receptor 2 (AstC-R2), which are homologous with mammalian somatostatin and its receptors^20^. AstC is released from EECs via a Target of Rapamycin (TOR)-dependent mechanism in response to nutrient deprivation and is required for the mobilization of stored lipids and carbohydrates to prevent hypoglycemia during periods of starvation. Gut-derived AstC signals the fasting state to AstC-R2 on the APCs and promotes food intake and energy mobilization by stimulating AKH secretion. These findings demonstrate that in response to fasting, AstC is released by EECs to coordinate systemic metabolic-stress responses, regulating energy homeostasis through the inter-organ regulation of AKH secretion. Our findings illustrate a mechanism by which gut-mediated nutrient sensing is required for the maintainance of organismal energy homeostasis during nutritionally challenging conditions. In response to fasting, gut-derived AstC enhances hypoglycemia-induced AKH secretion – conceptually similar to the role of incretin hormones in promoting insulin secretion when glucose is consumed – and AstC is key to maintaining metabolic homeostasis during nutrient deprived-states, and dysregulation of such gut-mediated signaling could contribute to metabolic disorders.

## Results

### Gut AstC signaling regulates organismal resistance to nutritional challenges

To identify intestinal hormones involved in nutrient sensing and in maintaining energy homeostasis during nutrient stress, we used RNAi to knock down secreted factors in the EECs of the adult *Drosophila* midgut and tested for changes in organismal starvation resistance (Fig. 1A). We restricted the RNAi effect to the adult stage using an EEC-specific GAL4 driver (*voilà-GAL4*) in combination with *Tubulin*-driven temperature-sensitive (TS) GAL80 (GAL80^TS^) expression, together referred to as *EEC-GAL4* (*EEC*>). We found that EEC knockdown of some factors including *Natalisin* resulted in sensitivity to starvation, while depleting others such as *Proctolin* and *AstC* enhanced starvation resistance. We focused our attention on AstC, since *UAS-AstC-RNAi* (*AstC-RNAi*)-mediated knockdown of this factor in the adult EECs (*EEC>AstC-RNAi*) caused a striking prolongation of starvation survival in females, but not in males, compared to *EEC>* controls (Fig. 1B and S1A). Sex-specific differences in metabolism are apparent in both humans and flies, but the underlying mechanisms are generally not well-understood. In *Drosophila*, male-specific fat metabolism is related to dimorphic expression of the lipase Brummer^21^, but the mechanisms underlying female-specific metabolic control remain to be characterized. To further confirm this phenotype, we used tissue-specific somatic CRISPR/Cas9-mediated gene editing to disrupt the *AstC* locus in the EECs of adult flies. For this purpose, we generated transgenic animals with UAS-inducible expression a pair of guide RNAs (gRNAs) targeting *AstC*, which was driven with *EEC>UAS-Cas9* (*EEC>Cas9*) containing GAL80^TS^ to restrict effects to the adult stage. Deletion of *AstC* in the EECs in the adult stage (*EEC>Cas9, AstC*^*KO*^) led to a strong increase in resistance to starvation in females compared to controls (*EEC>Cas9*), but not males (Fig. 1C and S1B), thus confirming the effect observed for RNAi-mediated knockdown. This phenotype suggests a post-developmental role of gut AstC in organismal responses to nutrient deprivation.

**Figure 1.**
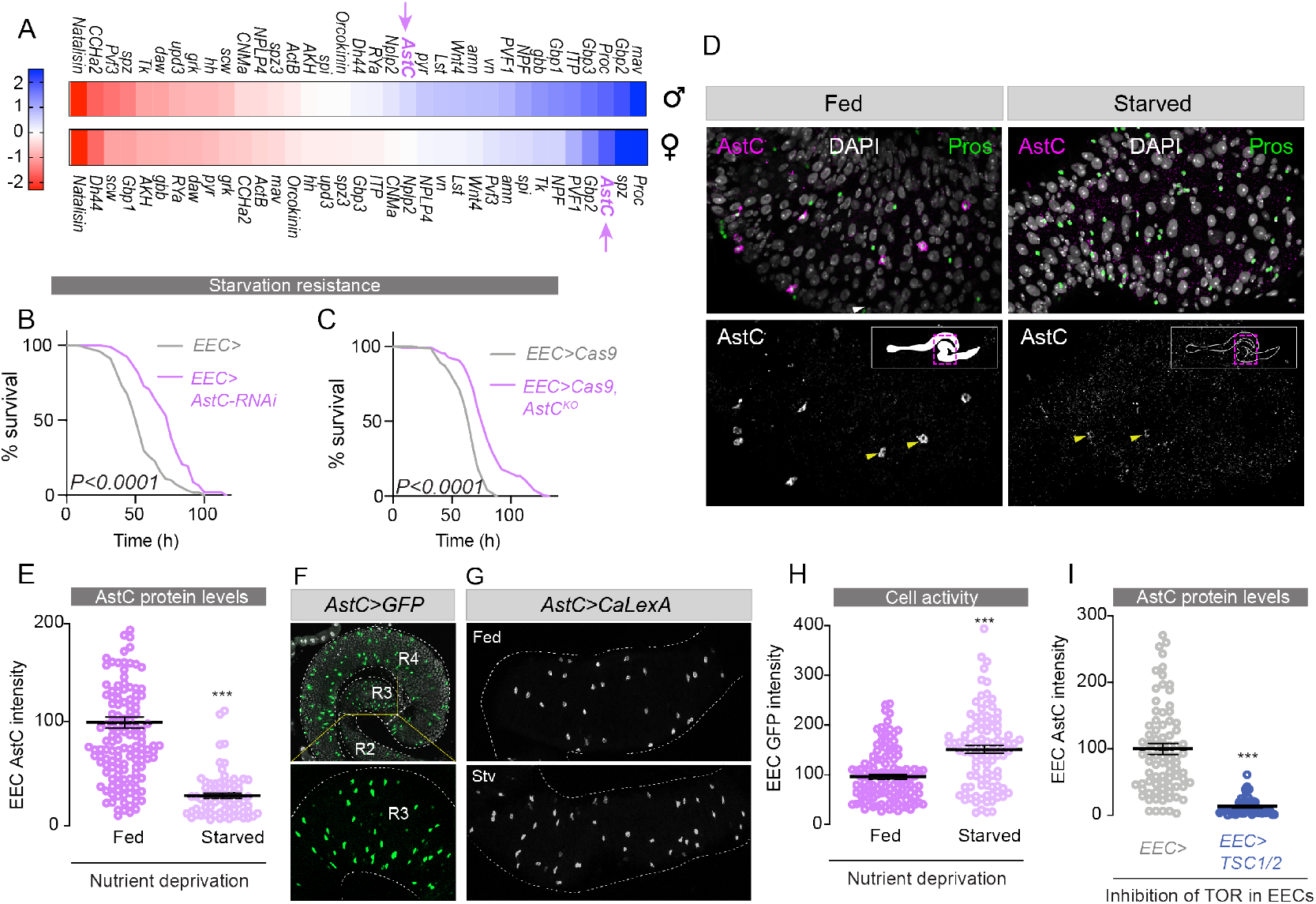
AstC secretion from the gut is nutrient-responsive and regulates organismal resistance to nutrient stress. **(A)** A screen of peptides expressed in the adult midgut identified peptides that shorten (red, left) or extend (blue, right) the survival of animals under starvation. Color scale indicates Z-score, the number of standard deviations the mean of each line falls from the mean of all lines. EEC-specific RNAi (**B**) or CRISPR-mediated knockout (**C**) of *AstC* prolongs female starvation survival (P<0.0001 by Kaplan-Meier log-rank test). **(D)** Starvation induces release of AstC (anti-AstC, magenta) from EECs (marked with anti-Pros, green). AstC staining is apparent in the EECs of fed females, whereas this staining is strongly reduced in starved animals, despite constant *AstC* expression (Fig. S1C). **(E)** Quantification of anti-AstC staining intensity shows a significant attenuation in peptide signal in the EECs of starved animals (results represent analysis of 9 guts). **(F)** AstC-expressing cells are abundant in the R3 and R4 regions of the female midgut (*AstC>GFP*, green; DAPI nuclear stain, white), and some AstC positive cells are also present in the R2 region. Bottom: an enlargement of the imaged R3 segment, where calcium-LexA signals were measured. AstC-expressing EECs in midgut segment R3 are activated by starvation, as indicated by the time-integrating calcium-LexA reporter system (*AstC>LexA::NFAT::VP16, LexAop-GFP*). GFP intensity, which reports recent calcium concentration, is increased in starved animals; these data (which represent analysis of guts from 6 animals) are quantified in **(H)**. **(I)** AstC release from EECs is inhibited by TOR activity. Acute expression of the TOR inhibitors *TSC1* and *TSC2* in EECs for six hours leads to a strong reduction in EEC AstC staining (results represent analysis of guts from at least 8 animals), even though *AstC* transcript levels increase with this manipulation (Fig. S1E). Error bars indicate standard error of the mean (SEM). ***, P<0.001.

### Nutrient-sensing in the EECs promotes release of AstC upon nutritional restriction

To be able to regulate organismal metabolism and food intake according to nutrient availability, the intestine must have the ability to couple nutrient-sensing with hormone secretion into circulation. To investigate whether EEC production or release of AstC is regulated by nutrients, we performed immunostaining of midguts from animals following feeding or nutrient deprivation. After 6 hours of starvation, AstC immunoreactivity was strongly reduced in EECs of adult female middle midguts (Fig. 1D and 1E). To exclude the possibility that this observation was due to reduced AstC production by the EECs, we performed gene expression analysis; *AstC* transcript levels were unchanged in adult midguts following 6 hours of starvation (Fig. S1C), suggesting that the depletion of AstC levels in the EECs in response to starvation can be attributed to increased release and not to decreased production. This indicates that AstC is released by EECs in response to nutrient restriction.

Activation of EECs results in elevation of their intracellular calcium levels^34^. We asked whether AstC-positive EECs of the adult female midgut are activated by nutrient restriction. First, we examined the expression pattern of *AstC-GAL4* (*AstC>*), a CRISPR knock-in of T2A::GAL4 at the C terminus of AstC^35^, in the female midgut by driving *GFP* expression to confirm *AstC-GAL4* (*AstC>*) expression in the EECs. *AstC>GFP* reporter activity is prominent in a large number of EECs of the middle (R3) and posterior (R4) regions and in a smaller number in the anterior R2 region (Fig. 1F). To measure the activity of AstC-positive EECs, we used the CaLexA reporter system to record intracellular calcium changes^28^. Starvation treatment for 6 hours increased calcium signaling in AstC-positive EECs as indicated by a larger number of GFP-positive AstC-expressing cells as well as increased GFP intensity in these cells (Fig. 1G, 1H, and S1D). This suggests that at least a subset of AstC-expressing EECs are activated when food is absent, consistent with the peptide’s being released during these conditions. We next sought to determine whether EECs sense and respond directly to levels of nutrients and relay availability systemically through release of AstC. We thus acutely inhibited in the EECs the activity of the Target of Rapamycin (TOR) pathway, a primary mediator of cell-autonomous nutrient sensing^8^, by overexpressing the TOR-inhibitory proteins *Tuberous Sclerosis Complex 1 and 2* (*TSC1/2*)^36^. This manipulation mimics nutrient deprivation in these cells in an otherwise nutrition-replete animal. After TOR inhibition was induced for 6 hours, AstC immunoreactivity was strongly reduced in the EECs, whereas intestinal *AstC* transcript levels were increased (Fig. 1I and S1E). These findings suggest that inhibition of TOR in the EECs activates the production and release of AstC into circulation by the intestine. Taken together our results suggest that intestinal TOR-mediated nutrient sensing in the EECs links low nutrient availability to the release of AstC.

### AstC signaling from the gut regulates metabolic adaptations to nutritional challenges

When animals, including *Drosophila*, encounter nutritional deprivation they must adapt by mobilizing stored energy for physiological and behavioral responses^1^. Their capacity to survive periods of nutrient scarcity is therefore directly correlated with their ability to make available stored body fat and carbohydrate resources. Whereas carbohydrates are stored mainly as glycogen and provide a fast-responding energy buffer, lipids are stored in the form of triglycerides (TAGs) in fat tissue and make up the main energy reserve. Thus, animals with increased levels of TAGs, or a reduced (but sufficient for survival) rate of their mobilization, are resistant to starvation. Given the effects of AstC on starvation survival, we investigated the physiological impact of *AstC* knockdown on flies’ ability to access their energy resources. Consistent with their increased resistance to starvation (Fig. 1B), we found that females lacking *AstC* in the EECs (*EEC>AstC-RNAi*) deplete both their stored glycogen and lipids more slowly than controls (*EEC>*). Whole-body glycogen levels were higher in females with *AstC* knockdown in the EECs after 15 hours’ starvation (Fig. 2A). Furthermore, females with EEC-specific *AstC* knockdown reduced their rate of TAG depletion, as indicated by higher whole-body TAG levels after 15 and 30 hours’ starvation compared to controls (Fig. 2B and 2C). When *AstC* was deleted in the adult EECs using CRISPR-mediated disruption of the gene (*EEC>Cas9, AstC*^*KO*^), females also depleted their TAG stores more slowly than controls (*EEC>Cas9*) upon starvation (Fig. S1F), further supporting the finding that lack of *AstC* in the EECs causes a deficiency in mobilizing stored energy. These effects of AstC on energy stores were not observed in males (Fig. S1G and S1H), consistent with female-specific effects on starvation, suggesting that gut-derived AstC plays a sexually dimorphic role in metabolic regulation.

**Figure 2.**
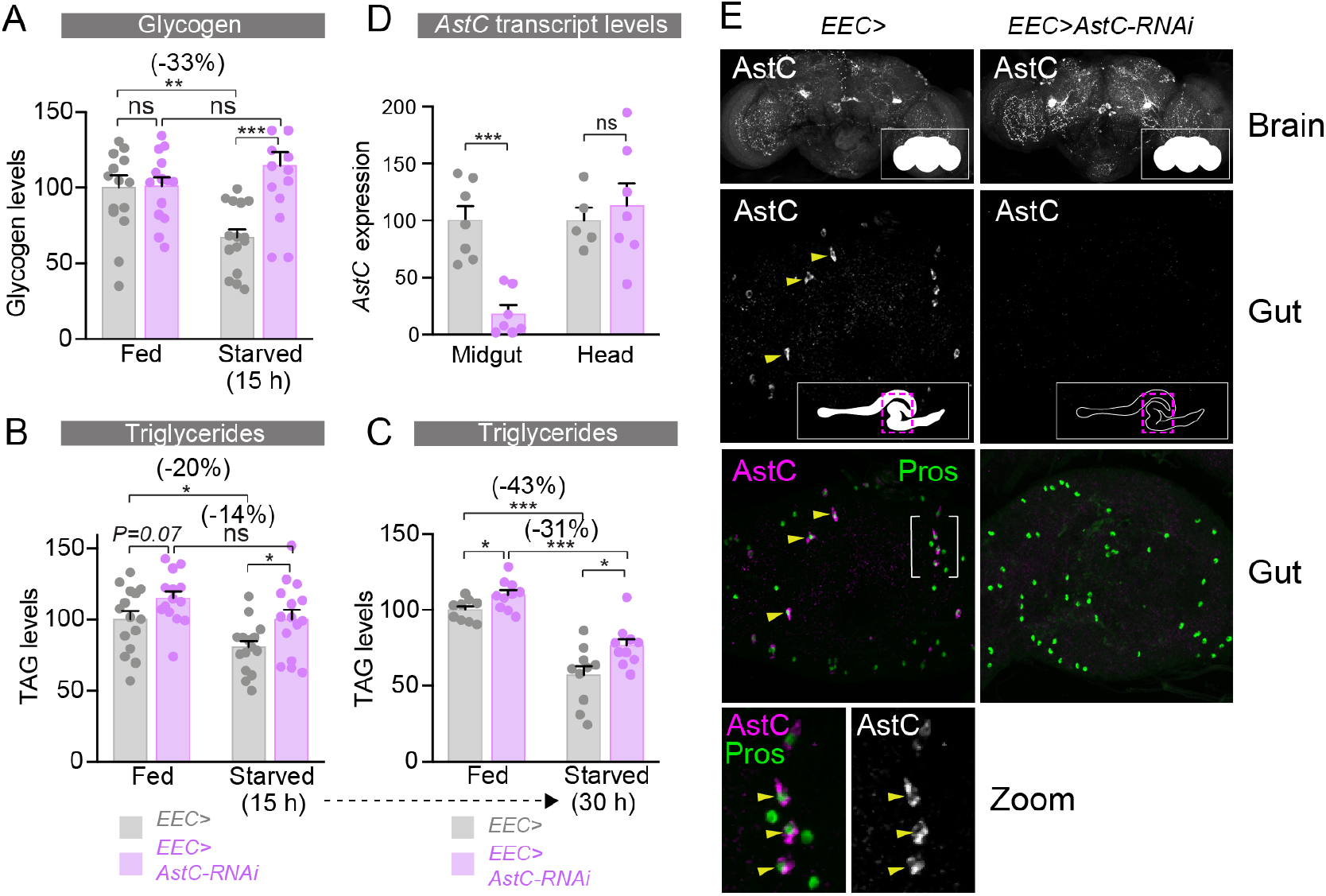
Gut-derived AstC is required for energy mobilization in response to nutritional challenges. **(A)** RNAi against *AstC* in the EECs prevents mobilization of stored glycogen during fifteen hours’ starvation. **(B)** *AstC* RNAi in the EECs reduces mobilization of triacylglycerides (TAGs) from adipose tissues during fifteen-hour starvation. **(C)** The mobilization-suppressing phenotype of *AstC* RNAi persists and is exacerbated with longer-term starvation (30 hours). **(D)** The genetic manipulation used to knock down *AstC* in the EECs (*EEC>AstC-RNAi*) very efficiently depletes midgut *AstC* transcript levels but does not alter neuronal expression of *AstC*. **(E)** Immunostaining of guts and brains shows likewise that brain AstC peptide distribution is not obviously reduced, whereas AstC expression (magenta) in the midgut EECs (marked in green with anti-Prospero) is strongly reduced. Below, a rotated and enlarged view of gut stain. Error bars indicate standard error of the mean (SEM). ns, non-significant; *, P<0.05; **, P<0.01; and ***, P<0.001.

Previous work has shown that AstC is also produced in a small set of neurons in the adult brain^22^. To attribute the observed phenotypes of *AstC* knockdown unambiguously to EEC depletion of AstC, we analyzed *AstC* transcript levels in dissected adult midguts and heads. We found a strong reduction of *AstC* transcripts in midguts from *EEC>AstC-RNAi* animals, while *AstC* expression was unaltered in the heads of these animals (Fig. 2D). We next performed anti-AstC immunostaining of adult brains and guts and found normal AstC signal levels in the brains of *EEC>AstC-RNAi* animals, whereas AstC was undetectable in the EECs of these animals (Fig. 2E). This demonstrates the tissue specificity of our RNAi knockdown and thus identifies the adult midgut EECs as the source of metabolism-governing AstC. Together, these results identify a nutrient-responsive role of EEC-derived AstC signaling that is necessary for metabolic homeostasis and adaptation to nutritional deprivation.

### AstC/AstC-R2 signaling regulates metabolic homeostasis through AKH

In *Drosophila*, nutrient homeostasis and energy mobilization are regulated by the antagonistic actions of two key metabolic hormones, the glucagon-like factor AKH and the insulin-like DILPs^1,8^. These signaling molecules regulate metabolic pathways that govern the storage and usage of energy, which is stored mainly as glycogen and TAGs in both flies and mammals. To assess the possible involvement of insulin and AKH signaling in mediating the effects of AstC on metabolic adaptation to nutritional challenges, we examined the expression of the two AstC receptors, AstC-R1 and AstC-R2, in the IPCs and APCs to gain insight into whether AstC might act directly on these two hormonal systems. To visualize receptor localization, we used transgenic lines in which CRISPR-mediated insertion of *T2A::GAL4* into the native *AstC-R1* and *AstC-R2* loci allows *UAS-GFP* reporter-based analysis of AstC-R1 and AstC-R2 expression patterns^23^. We found that AstC-R2, but not -R1, is expressed in the APCs, as indicated by co-localization with anti-AKH immunofluorescence (Fig. 3A and S2A). Similarly, AstC-R2, but not -R1, is expressed in the adult IPCs, as indicated by overlap with anti-DILP2 signal (Fig. S2B).

**Figure 3.**
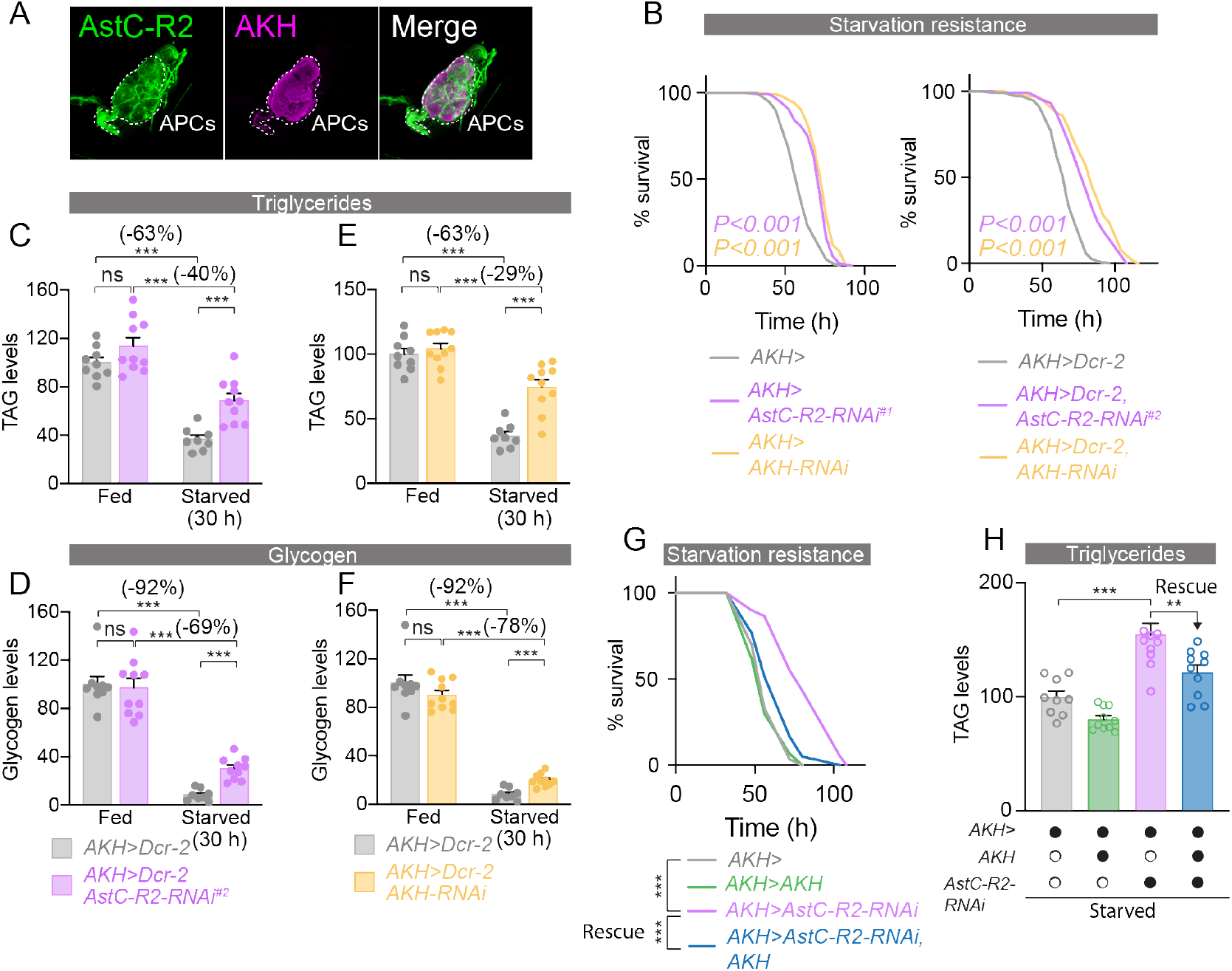
AstC acts via AstC-R2 in the APCs to regulate AKH signaling, which mediates the metabolic effects of AstC signaling. **(A)** *AstC-R2::2A::GAL4*-driven *UAS-GFP* expression (left, green) is apparent in the AKH-producing cells (APCs) of the CC, marked with anti-AKH (magenta). **(B)** Knockdown of *AstC-R2* in the APCs phenocopies the starvation-survival effect of *AstC* knockdown in the gut; knockdown of *AKH* in these cells also reproduces this phenotype. **(C and D)** APC-specific knockdown of *AstC-R2* reduces the consumption of TAGs **(C)** and glycogen **(D**) during starvation. **(E and F)** Likewise, APC knockdown of *AKH* itself induces similar reductions in starvation-induced mobilization of TAGs **(E)** and glycogen **(F). (G)** The effects of AstC signaling are mediated by AKH: overexpressing *AKH* restores normal starvation sensitivity in animals with APC-specific knockdown of *AstC-R2*. **(H)** Likewise, overexpression of *AKH* in animals with APC-specific knockdown of *AstC-R2* rescues their deficiency in lipid mobilization during starvation. Error bars indicate standard error of the mean (SEM). ns, non-significant; and ***, P<0.001.

To assess the potential contribution of AKH and DILP signaling to the metabolic phenotypes associated with *AstC* loss in the EECs, we used RNAi to knock down *AstC-R2* in the APCs or IPCs. Since loss of AKH signaling, like loss of gut-derived AstC, leads to increased resistance to starvation, we hypothesized that AKH might mediate the metabolic effects of AstC. To test this, we knocked *AstC-R2* down in the adult APCs and measured organismal resistance to nutritional deprivation. APC-specific knockdown of *AstC-R2* using *AKH-GAL4*, restricted to the adult stage with GAL80^TS^ (together referred to as *AKH>*), led to a marked increase in resistance to starvation in females with one *UAS-AstC-R2-RNAi* line (referred to as *AstC-R2-RNAi*^*#1*^), but not males, phenocopying the effects of *AstC* knockdown in the EECs (Fig. 3B and S2C). In contrast, adult-specific knockdown of *AstC-R1* in the APCs did not prolong survival under starvation stress (Fig. S2D), ruling out a large contribution of this receptor to AstC-mediated effects on the APCs. We confirmed the starvation-resistance phenotypes of AstC-R2 knockdown in the APCs using another independent RNAi line, whose effectiveness was increased through the co-expression of the RNA-processing enzyme Dicer 2 (Dcr-2) (Fig. 3B), as well as adult-APC-specific CRISPR/Cas9-mediated knockout of *AstC-R2* (Fig. S2E) driven by *AKH>GAL80*^*TS*^, *Cas9*. This indicates that the phenotype is caused specifically by loss of *AstC-R2* in the APCs and suggests that loss of AstC signaling in the APCs contributes to the increased survival of starved animals with EEC-specific AstC depletion.

The metabolic stress experienced during nutritional deprivation elicits hormonal responses that mobilize stored resources, redistribute energy towards essential physiological functions, and initiate food-seeking behaviors^1^. AKH secretion induced by hypoglycemia promotes mobilization of TAG from the fat body. AKH-deficient animals are therefore more resistant to starvation due to their slower mobilization of TAG from the fat body^24^. We also assessed resistance to starvation in animals with APC-specific RNAi against *AKH* (Fig. 3B), confirming previous work showing that loss of AKH signaling leads to increased starvation resistance^24^, thereby mimicking disruption of AstC signaling. Taken together, our results show that loss of *AstC* in the EECs or of *AstC-R2* in the APCs leads to prolonged survival under starvation, implying that AstC signals to the APCs during nutritional restriction to promote systemic energy mobilization. To determine whether AstC signaling acts on the APCs to promote energy mobilization by peripheral tissues, we analyzed whole-body TAG and glycogen levels of animals with *AstC-R2* depletion in the APCs to assess their energy stores. Adult-restricted *AstC-R2* knockdown in the APCs led to higher whole-body TAG levels after starvation in females compared to controls (Fig. S2F), similar to the effects observed with knockdown of *AstC* in the EECs, indicating a post-developmental role of AstC-R2 in the regulation of APC activity. Independent validation of these results was obtained with the second *AstC-R2-RNAi* line in combination with *Dcr-2* co-expression, which led to an even more severe metabolic phenotype in which animals depleted their stored TAGs and glycogen significantly more slowly in response to starvation (Fig. 3C and 3D). The effect of *AstC-R2* RNAi was similar to the phenotype observed in animals with loss of *AKH* signaling by APC-specific knockdown of *AKH*, which led to a reduced rate of TAGs and glycogen depletion during nutritional deprivation (Fig. 3E, 3F, and S2G). Together these findings suggest that AstC promotes lipid and carbohydrate mobilization by direct action on the APCs to promote the release of AKH during nutritional restriction.

To determine whether AKH signaling mediates the effects of AstC on resistance to starvation and metabolic stress, we assessed whether simultaneous *AKH* overexpression might rescue these phenotypes. Co-expression of *AKH* along with *AstC-R2-RNAi* was sufficient to revert the starvation-resistance and energy-mobilization phenotypes caused by *AstC-R2* knockdown alone in the APCs (Fig. 3G and 3H), suggesting that impaired AKH signaling is the primary cause of the metabolic-stress phenotypes observed in *AstC*-deficient animals. Together, our findings suggest that communication between gut and APCs is mediated by AstC, which acts on AstC-R2 on the APCs to promote AKH secretion. This indicates that signaling between nutrient-responsive AstC-expressing EECs and the neuroendocrine CC is required for potentiation of AKH release during nutritional restriction.

Because AstC-R2 is also expressed in the adult female IPCs (Fig. S2B), we also investigated the possibility that DILP signaling might mediate some of the effects of gut-derived AstC on the metabolic response to starvation. Knockdown of *AstC-R2* in the IPCs had no effect on starvation-induced depletion of TAG and glycogen in females (Fig. S3A and S3B), indicating that energy mobilization promoted by EEC-derived AstC is not mediated through effects on IPC secretion of DILPs. Furthermore, we found no difference in the levels of DILP2 or -3 peptides in the IPCs after starvation in females with *AstC* knockdown in the EECs or IPC knockdown of *AstC-R2* (Fig. S3C and S3D), indicating that gut AstC does not strongly influence DILP secretion upon nutritional restriction. We also analyzed whether *AstC* loss in the EECs affects systemic insulin signaling by measuring levels of phosphorylated AKT (pAKT), a downstream target of insulin signaling^8^. Adult females with EEC-specific knockdown of *AstC* showed no change in whole-body pAKT levels compared to controls in fed conditions or after 6-hour starvation (Fig. S3E). Collectively, these data indicate that the stress resistance and metabolic phenotypes associated with EEC-specific depletion of AstC are mediated by AKH signaling and are not due to effects on DILP signaling.

### Gut-derived AstC promotes AKH secretion in response to nutritional challenges and prevents hypoglycemia

AKH released in response to nutrient stress promotes mobilization of resources from the fat body via its receptor AKHR to maintain energy homeostasis^10^. To further study the function of AstC in controlling AKH signaling, we examined the binding of fluorescently labelled AstC to the APCs. Using this ligand-receptor binding assay^25^, we found that AstC binds to the APCs in females, as indicated by co-localization of labelled AstC with *AKH>-*driven GFP fluorescence (Fig. 4A). Furthermore, this binding of AstC was abrogated by knockdown of *AstC-R2* in the APCs, suggesting that AstC binds to AstC-R2 on the APCs.

**Figure 4.**
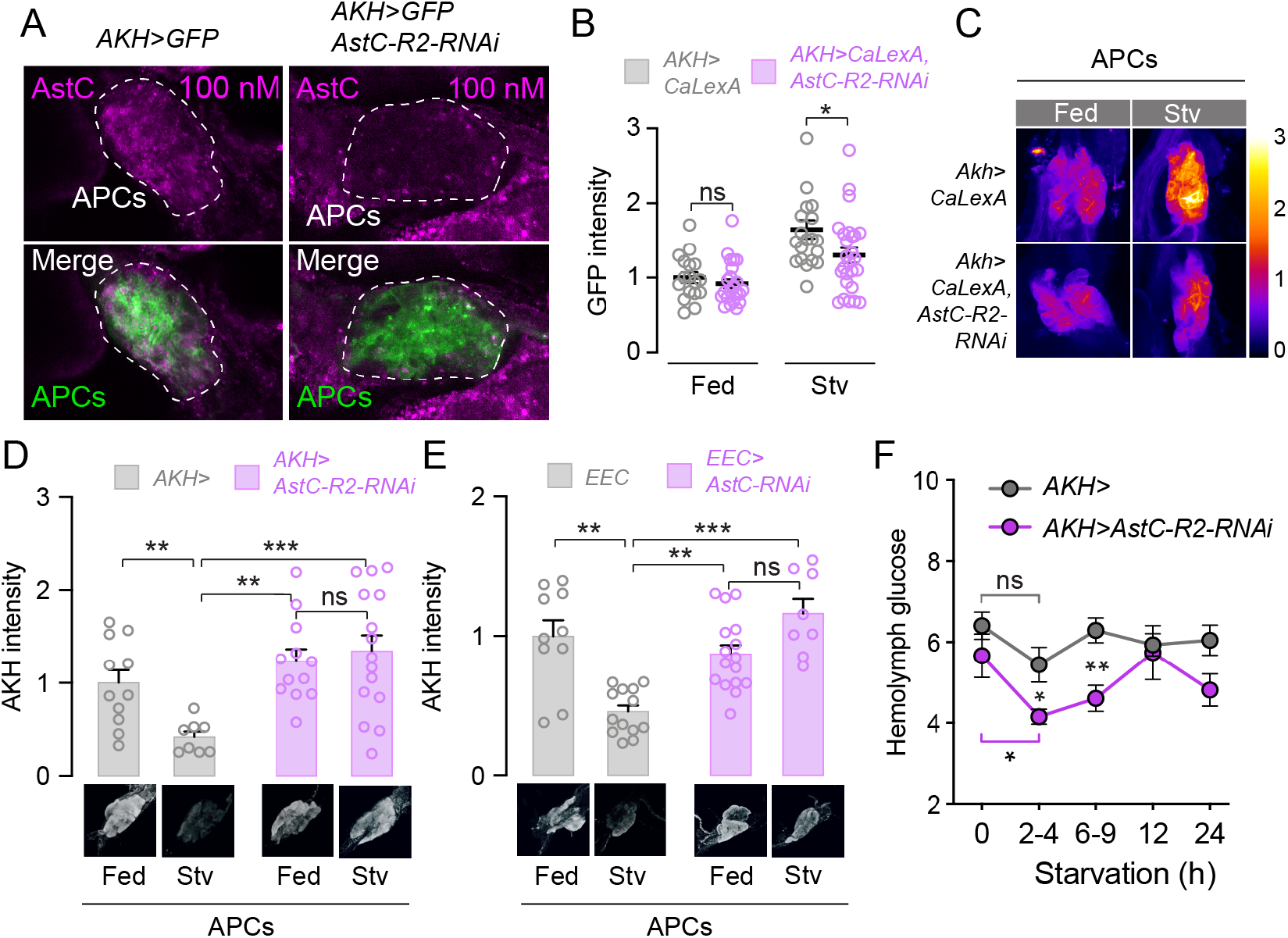
AstC acts on the APCs to promote AKH release and thereby maintain glycemic levels. **(A)** Fluorescently labeled AstC peptide (magenta) binds to the APCs (marked by *AKH>GFP*, green) of normal animals, whereas staining is much weaker in APCs expressing *AstC-R2*-*RNAi*. **(B)** AstC potentiates the starvation-induced activation of the APCs. Calcium-induced CaLexA-GFP signal intensity in the APCs is strongly increased by starvation in control animals (gray), whereas this increase is attenuated by RNAi against *AstC-R2*. **(C)** Illustrative images of calcium-induced GFP expression in the APCs. **(D)** AstC-R2 signaling in the APCs is required for starvation-induced AKH release. Immunostaining of the processed AKH peptide shows that starvation induces peptide release in control animals, whereas the intracellular concentration of AKH is not affected by starvation if *AstC-R2* is knocked down in the APCs. See also Fig. S4A. **(E)** Likewise, AstC from the gut is required for starvation-induced AKH release. Illustrative anti-AKH images of the APCs are shown below each bar. **(F)** AstC signaling in the APCs is required for the maintenance of circulating sugar levels in the hemolymph during starvation. Error bars indicate standard error of the mean (SEM). ns, non-significant; *, P<0.05; **, P<0.01; and ***, P<0.001.

Pancreatic *α* cells respond to hypoglycemia with an increase in their intracellular calcium levels, which promotes glucagon secretion^26^. In *Drosophil*a, decreased extracellular sugar concentration likewise induces increased calcium signaling in explanted APCs, suggesting that similar mechanisms regulate secretion of glucagon and AKH^27^. To test whether nutrient deprivation enhances calcium signaling in the APCs, we used the CaLexA system to report intracellular calcium concentration. Using this approach, we found that the APCs show elevated calcium-reporter activity after *in vivo* starvation treatment (Fig. 4B and 4C), indicating that APCs behave similarly to pancreatic *α*-cells in response to nutrient deprivation. By contrast, starvation-induced APC calcium activity was strongly attenuated by knockdown of *AstC-R2*, demonstrating that a large part of the APC response to nutrient deprivation requires AstC signaling. Taken together these findings suggest that activation of the APCs by gut-derived AstC elevates intracellular calcium levels, inducing AKH secretion and thus promoting the mobilization of stored lipids and sugars from peripheral tissues.

To assess whether AstC regulates AKH production and secretion upon nutritional challenges, we first measured *AKH* transcript levels. Consistent with starvation-induced upregulation of AKH signaling, we found that *AKH* expression was increased in females upon starvation (Fig. S4A), indicating increased AKH production. This effect was abolished by knockdown of *AstC-R2* in the APCs, suggesting that AstC signaling within the APCs is required for *AKH* upregulation upon nutrient deprivation. We then asked whether AKH release during nutrient stress also requires gut AstC signaling. To test whether AstC signaling regulates the secretory activity of the APCs, we analyzed their intracellular AKH protein levels upon starvation. Starvation treatment for 6 hours led to a significant drop in AKH staining levels in control females, whereas no differences in intracellular AKH levels between fed and starved conditions were observed in animals with *AstC-R2* knockdown in the APCs (Fig. 4D). This indicates that AstC-R2 is required in the APCs for starvation-induced release of AKH. We then asked whether AstC released from the gut is required for the AKH release observed upon starvation. EEC knockdown of *AstC* completely abolished starvation-induced AKH release, as indicated by unchanged intracellular AKH levels (Fig. 4E), suggesting that gut AstC in a non-cell-autonomous manner promotes AKH secretion in response to metabolic challenges. Furthermore, receptor upregulation is a common mechanism by which cells can increase sensitivity in response to low abundance of ligand. We observed increased *AkhR* upregulation in response to starvation in females with EEC-specific loss of *AstC*, consistent with low AKH signaling (Fig. S4B). Since nutrient deprivation increases *AKH* gene expression as described above (Fig. S4A), these findings collectively suggest that gut-derived AstC positively modulates AKH production and release during nutrient restriction.

Glucagon acts to prevent hypoglycemia^29^, and impaired AKH-induced breakdown of energy stores could result in hypoglycemia upon nutritional challenge. We therefore examined animals’ ability to maintain normal circulating sugar levels during fasting. To assess whether AstC-mediated AKH secretion has an impact on circulating sugar, we collected hemolymph before and after starvation. *AstC-R2* knockdown in the APCs led to fasting-induced hypoglycemia (Fig. 4F), consistent with an impairment of AKH-mediated energy mobilization. Control females maintained relatively constant hemolymph glucose without significant changes during 24 hours of fasting, while hemolymph glucose dropped significantly within a few hours of fasting when *AstC-R2* function was depleted in the APCs. Consequently, females with APC-specific *AstC-R2* knockdown exhibited reduced circulating sugar levels, indicating they were in a hypoglycemic state for several hours following starvation. These findings together indicate that AstC released by the endocrine cells of the gut in response to nutrient deprivation acts on the APCs to promote AKH release and thus to promote energy mobilization to prevent hypoglycemia.

### Gut-derived AstC promotes food intake via AKH signaling

Homeostatic control of energy balance requires the coordination of food intake and energy expenditure^1^. Metabolic processes are therefore intimately coupled with regulation of feeding to maintain a constant internal state. To investigate whether AstC signaling coordinates metabolic and behavioral responses to nutrient deprivation, we monitored food intake in animals with adult-restricted EEC-specific *AstC* knockdown. *AstC* loss in the EECs caused a significant decrease in short-term (30-minute) food intake measured by dye assay in females (Fig. 5A), whereas food intake was not reduced in males (Fig. S5). To further understand the endocrine mechanisms by which *AstC* deficiency causes hypophagia, we tested whether AstC regulates food intake through its action in the APCs, using adult-restricted knockdowns. Using the short-term dye-feeding assay, similar results were obtained for females when *AstC-R2* or *AKH* were knocked down in the APCs as when *AstC* was depleted in the in EECs (Fig. 5A). Next, we quantified long-term food consumption using the CAFE assay^30^ to monitor food intake over 24-hour periods. Consistently, long-term food consumption was reduced in females when *AstC* was knocked down in the EECs or when *AstC-R2* or *AKH* itself were knocked down in the APCs (Fig. 5A). We then tested the impact of AstC signaling on feeding regulation using the FLIC (fly liquid-food interaction counter) system, which enables analysis of *Drosophila* feeding behaviors^31^. We found that adult-specific knockdown of *AstC* in the EECs reduced the number of tasting and feeding events, while increasing the time interval between feeding events over 24 hours (Fig. 5B). Thus, despite having similar TAG and glycogen levels in the fed state, as shown above, females with reduced AstC signaling are hypophagic. This shows that the metabolic phenotype is not caused by increased food intake or impaired nutrient absorption. Together our results indicate that in response to nutrient deprivation, AstC signaling from the gut promotes hunger and food consumption through regulation of AKH, consistent with previous findings showing that AKH is an orexigenic peptide^32,33^.

**Figure 5.**
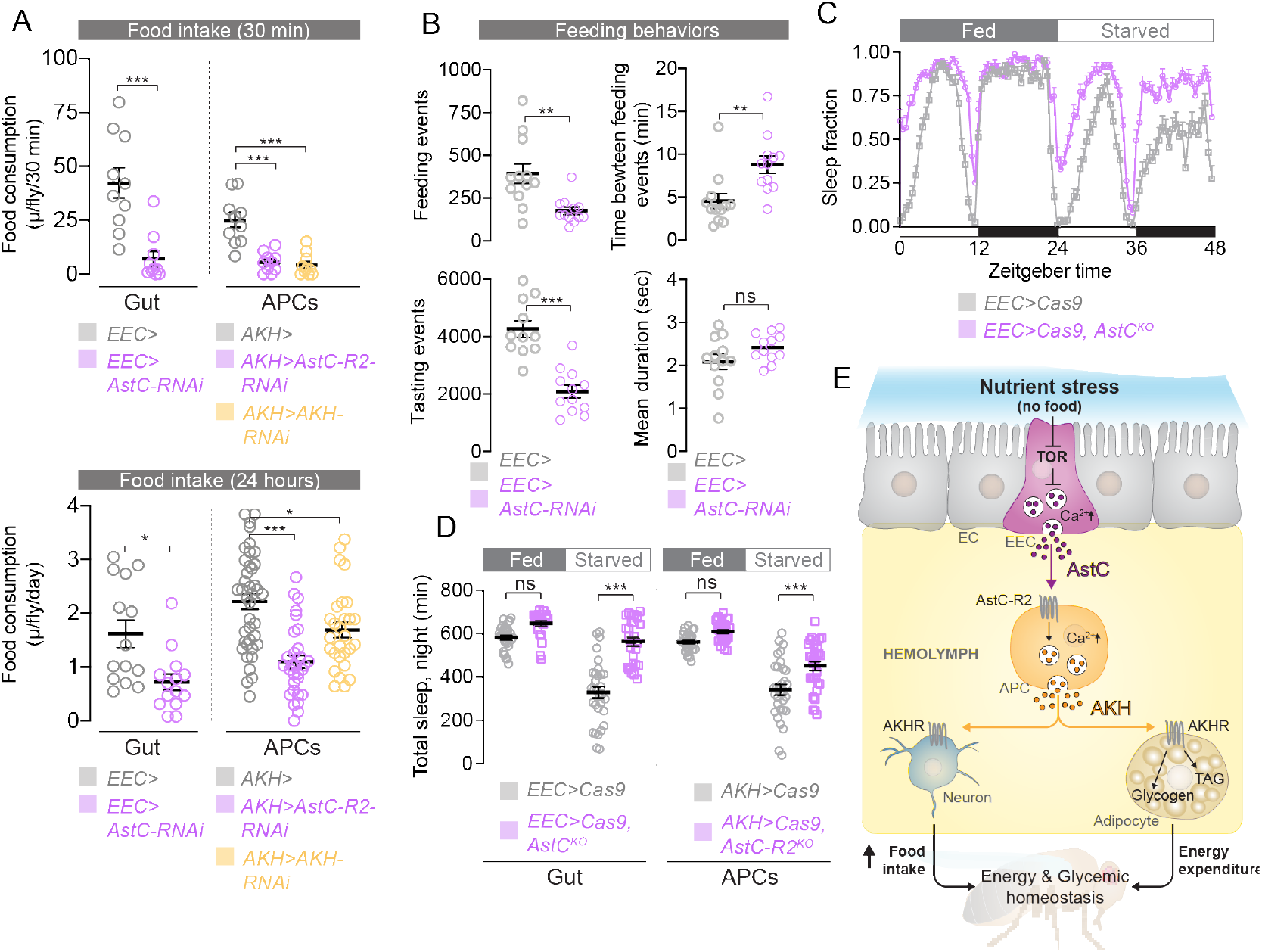
AstC signaling promotes food-seeking behaviors. **(A)** Top panel: short-term dye feeding assays indicate that knockdown of *AstC* in the EECs (magenta) or of *AstC-R2* or *AKH* in the APCs (orange) reduces food consumption relative to controls (gray). Bottom panel: longer-term CAFE feeding shows a similar effect, with knockdown of either *AstC* in the gut or of *AstC-R2* or *AKH* in the APCs leading to reduced consumption. **(B)** Long-term monitoring of feeding behaviors using the FLIC system indicates that knockdown of *AstC* in the midgut EECs (magenta) reduces “feeding” and “tasting” events and increases the interval between feeding events, whereas the duration of individual events is unaltered. **(C)** Monitoring of locomotor activity using the DAM system indicates that knockout of *AstC* in the EECs (magenta) does not strongly affect sleep architecture compared to controls (gray) during fed conditions (hours 0 through 24); during starvation (hours 24 through 48), however, wild-types exhibit a strong sleep-suppression phenotype that is strongly attenuated in CRISPR knockout animals. **(D)** Quantification of total nighttime sleep duration shows that starvation induces a strong reduction in nighttime sleep in control animals (gray); this effect is attenuated in animals in which *AstC* function in the EECs or *ActC-R2* function in the APCs have been disrupted by CRISPR-mediated gene deletion (magenta). **(E)** A model of AstC-mediated gut nutritional signaling. Reduced nutritional levels are sensed by TOR in the enteroendocrine cells (EECs); reduced TOR activity disinhibits EEC release of AstC into the hemolymph, which carries the peptide to its target tissues, including the AKH-producing cells (APCs) of the corpora cardiaca (CC). Although the APCs can autonomously sense circulating sugar levels, they also require AstC input via AstC-R2 to potentiate their activity during starvation, increasing intracellular Ca^++^ levels and promoting AKH expression and release into the hemolymph. Circulating AKH acts via AkhR (1) in the fat body, inducing the breakdown and mobilization of stored energy, and (2) in feeding- and arousal-modulating cells of the nervous system to suppress sleep, upregulate locomotion, and promote feeding behaviors. Through these effects, gut-derived AstC promotes behavioral and physiological responses to starvation that tend to promote energetic and nutritional homeostasis. Error bars indicate standard error of the mean (SEM). ns, non-significant; *, P<0.05; **, P<0.01; and ***, P<0.001.

Organisms also respond to nutritional challenges by alterating their food-seeking behavior. In *Drosophila*, starvation induces hyperactivity and suppresses sleep, presumably to promote food-seeking activities. Since AKH is required for starvation-induced hyperactivity and sleep inhibition, we investigated whether AstC is important for sleep responses to food deprivation. We monitored sleep under normal feeding and for 24 hours following nutrient deprivation in animals with *AstC* knockout in the EECs. Under fed conditions, control and knockout animals displayed similar sleep patterns (Fig 5C and 5D); under starvation, however, while controls exhibited a strong reduction in sleep, females with EEC-specific *AstC* deletion showed almost no sleep loss (Fig. 5C and 5D). Next, we tested whether AstC regulates sleep loss during starvation via AstC-R2 in the APCs. As seen with knockout of *AstC* in the EECs, females with *AstC-R2* knockout in the APCs resisted, at least partially, the starvation-induced sleep loss (Fig. 5D). These results indicate that AstC, via the APCs, regulates behavioral responses to starvation in order to initiate food-seeking. Combined, our findings show that gut-derived AstC controls feeding by AstC-R2-mediated effects in the APCs that potentiate hypoglycemia-induced AKH secretion. This indicates that under conditions of nutrient restriction, AstC signaling promotes hunger and food-seeking behaviors and therefore enhances food consumption to coordinate the organism’s homeostatic regulation of food intake and energy usage.

## Discussion

### Gut-derived AstC regulates metabolism and food intake to maintain energy homeostasis under nutrient-stress conditions

The gut has emerged as a central signal-integrating regulator of whole-body energy homeostasis. Despite the evolutionary divergence of flies and humans, the guts of these animals exhibit physiological and structural similarity, with the *Drosophila* midgut functioning analogously to the human small intestine^6,7^. Work in mammals and *Drosophila* has shown that the gut integrates nutritional and microbiotic cues and converts these into hormonal signals that relay nutritional information systemically to other tissues. However, our understanding of the cellular and molecular basis of nutrient sensing in the EECs, as well as the hormones these cells release, is limited. We conducted a screen of EEC-expressed peptide hormones for effects on starvation survival. Among our strongest survival-prolonging hits was AstC, one of the most highly and broadly expressed peptides in the *Drosophila* midgut^37,38^. Despite its abundance, the function of gut-derived AstC has remained unknown. Here, we show that certain midgut EECs respond to nutrient deprivation by secreting AstC, which ultimately acts to induce energy mobilization and to promote food-seeking behaviors (Fig. 5E). Retention of AstC under fed conditions requires the activity of the intracellular nutrient sensor TOR, which is promoted by nutrients, particularly amino acids^8^. AstC acts directly upon the APCs, which express the receptor AstC-R2, to promote AKH release. Playing a role analogous to that of mammalian glucagon, AKH acts on its receptor AkhR in adipose tissue to induce breakdown of energy stores, thereby protecting against deleterious drops in hemolymph sugar levels leading hypoglycemia during fasting or periods with rapid energy expenditure^24^.

Nutrient stress also modulates feeding decisions and behaviors to optimize survival. Nutritional deprivation increases locomotor activity and suppresses sleep in animals as an adaptive response to promote food-seeking behaviors^24,39^. AKH mediates this starvation-induced behavioral arousal response through its action on a small number of octopaminergic neurons in the brain that express AkhR^11^. Like human glucagon, AKH also has direct orexigenic effects, acting through four interoceptive AkhR-expressing neurons in the suboesophageal zone to promote consumption^33^. Our findings show that gut-derived AstC signaling is required for starvation-induced sleep suppression and feeding increase via actions that are mediated, at least in part, by AstC-R2 in the APCs. Together our results suggest that starvation induces AstC release from EECs into the hemolymph to increase appetite, arousal, and food-seeking behaviors, along with breakdown of energy stores, through regulation of AKH (Fig. 5E). Thus, AstC coordinates behavioral and metabolic responses which allows organismal adaptations to nutritionally stressful environmental conditions.

In contrast to the regulation of insulin release, which is well studied, the mechanisms governing glucagon secretion are poorly defined^40^ and have recently become a focus of intensive research due to their relevance to diabetes. Likewise, in the fly, the regulation of insulin secretion has been examined intensively, whereas the mechanisms by which AKH secretion is controlled are poorly understood, although the APCs, like mammalian pancreatic *α*-cells, are believed to be competent to sense circulating sugar levels^27^. Our present findings show that animals lacking *AstC* in the EECs, or its receptor *AstC-R2* in the APCs, fail to regulate their mobilization of stored energy properly during nutrient deprivation and thus deplete their stores of TAGs and glycogen more slowly than normal; as a consequence, these animals exhibit prolonged starvation survival, at the expense of becoming hypoglycemic. Thus, AstC signaling is required for AKH secretion in response to fasting to mobilize energy and avoid hypoglycemia. Paradoxical glucagon secretion under hyperglycemia is frequently observed in diabetic patients^41^. Thus, dysregulation of signals that act similarly to AstC, promoting glucagon-like signaling, may contribute to hyperglycemia in diabetes and therefore be of therapeutic relevance.

*Drosophila* AstC-R2 is orthologous with mammalian somatostatin receptors^20^, and the processed AstC peptide shares structural features with somatostatin, both being cyclized via disulfide bridges. Somatostatin (as its name suggests) is known for its regulatory effects on growth hormone, although it has recently also received attention for its role in the regulation of glucagon secretion^42^. Two active forms of the somatostatin peptide, which elicit partially overlapping responses, are produced by different tissues. Like AstC, which is produced by a small number of neurons in the *Drosophila* brain and is also highly expressed in the fly intestine, mammalian somatostatin is produced by the hypothalamus, but the major source of circulating peptide is the gastrointestinal tract. Endocrine cells of the gut produce the long somatostatin-28 isoform^43,44^, and the *δ* (delta) cells of the pancreas synthesize the short somatostatin-14 isoform, which has a strong paracrine intra-islet regulatory effect on glucagon secretion. However, less is understood about whether systemic somatostatin signaling from the intestine might signal to the pancreas and exert a role there in the regulation of glucagon secretion. Our results demonstrate that in *Drosophila*, the EECs release AstC systemically, which signals directly to the APCs and regulates AKH secretion, suggesting the possibility that mammalian gut-derived somatostatin may play a similar role in regulating glucagon secretion. We propose that AstC/somatostatin and AKH/glucagon define an evolutionarily conserved endocrine module regulating homeostatic control of food intake and metabolism.

In addition to processing ingested food, the intestine functions as a barrier between the external and internal environments that plays a key role in immunity, and the gut microbiome plays important roles in host physiology and health^45^. The mechanisms by which gut bacteria affect host physiology are believed to include the sensing of microbial metabolites by EECs, which respond by releasing hormones that act as cytokines^46^. Interestingly, AstC signaling mediated by AstC-R2 is also involved in *Drosophila* inflammatory and immune responses^47^. Other nutrient-sensitive gut-derived hormones appear to regulate both of these pathways as well, such as Bursicon signaling through its receptor Lgr2/Rickets^18,48^, and the interconnected effects of gut signaling on metabolism and immunity will be an interesting area for future exploration.

Males and females differ in their metabolic physiology and susceptibility to disease^49^. These differences are likely associated with distinct male and female reproductive strategies and lifestyles. In both mammals and *Drosophila*, females store more lipids than males do^21,50^. Female flies also consume more food than males^1^ and have a higher demand for dietary amino acids^51^, which are required to sustain metabolically demanding oogenesis, whereas reproduction is less metabolically costly for males. Our finding of female-specific AstC function raises the possibility that this peptide may affect some of these previously noted female-specific metabolic and behavioral adaptations, contributing to greater female reproductive success. Females also must promote the developmental success of their offspring, which require dietary proteins for their development. Females may therefore respond to a lack of such nutrients by initiating food-seeking behaviors in order to locate rich sites in which to deposit their eggs. It will be of interest for future studies to further explore the female-specific function of AstC, including whether AstC is linked to behaviors and physiology associated with female reproduction. Our observations of sex-specific differences are perfectly in line with recent work showing that sexual dimorphism in TAG metabolism in *Drosophila* governs starvation resistance in males and females^21^. In males, this sex-specific difference depends on the lipase Brummer that promotes lipolysis of stored TAG in response to starvation. Our results indicate a role of AstC in regulation of sex-specific differences that underlie female-specific TAG metabolism in response to nutritional challenges, which may be important for a better understanding of sex differences in metabolism and disease therapies.

### Gut-mediated nutrient sensing and signaling and their implications for metabolic diseases

Balance between food intake and energy expenditure is necessary for the maintenance of energy homeostasis, and imbalance in these processes can lead to obesity or wasting. Organismal energy balance therefore requires the ability to sense food cues and to regulate feeding and metabolism appropriately in response^1^. Nutrient-sensing pathways and the regulation of metabolism and feeding have become the focus of intense research, as these are the keys to controlling energy balance and are often dysregulated in human metabolic disorders^52^. Intestinal nutrient sensing has emerged as a major endocrine regulator of body-wide metabolism, food intake, and blood-glucose control, and targeting intestinal nutrient sensing and downstream gut-derived hormones holds great promise for the treatment of diabetes and obesity. The ability of the gut to sense nutrients is critical for regulating the release of gut hormones involved in appetite and metabolic control. Our work shows that gut AstC release is regulated by TOR-dependent nutrient sensing in the EECs, also indicating that the APCs sense nutrient status indirectly via the EECs and AstC. TOR is activated by amino acids, but glucose also promotes TOR activity by inhibiting the TOR-inhibitory energy sensor AMPK and thus disinhibiting TOR^53^. Thus, deprivation of amino acids or glucose inhibits TOR, which is a primary mechanism by which cells and organisms sense scarcity and induce adaptive growth and metabolic responses to dietary restriction. We now provide evidence that intestinal nutrient sensing via TOR controls body-wide metabolism and feeding through the release of AstC, which promotes food intake and energy expenditure.

In mammals, glucose sensing in the EECs is a main mechanism by which insulin secretion is regulated, since the combined actions of the incretin hormones GLP-1 and GIP, both released from EECs upon the ingestion of glucose, potentiate hyperglycemia-induced insulin secretion^54^. Incretin effects are impaired in diabetic patients, and clinical use of GLP-1-agonist therapies has recently become a widely used strategy for the treatment of diabetes and obesity. In these metabolic disorders, impaired intestinal nutrient-sensing and incretin signaling leads to a failure to maintain glycemic homeostasis and regulate food intake^55^. We show here that AstC is a hormone secreted from intestinal EECs that induces AKH secretion, providing the first evidence of a gut hormone with potential incretin-like properties that acts on glucagon-like signaling systems. Taken together our findings identify an endocrine pathway that regulates homeostatic control of food intake and energy metabolism. Identification of hormonal factors such as AstC that control food intake and energy metabolism may be exploited to control appetite and body weight and help pave the way for development of new strategies for the treatment of diabetes and obesity.

## Methods

### *Drosophila* stocks and husbandry

Flies were maintained on a standard cornmeal medium (containing 82 g/L cornmeal, 60 g/L sucrose, 34 g/L yeast, 8 g/L agar, 4.8 mL/L propionic acid, and 1.6 g/L methyl-4-hydroxybenzoate) at 25 °C and 60% relative humidity under normal photoperiod (12 h light: 12 h dark). Genotypes with temperature-sensitive GAL80 (*Tub-GAL80*^*TS*^) were raised through eclosion at 18 °C. Newly eclosed flies were kept at 18 °C for 3-4 days more prior to separation by sex (15 flies per vial) and transferred to 29 °C for five days to induce UAS expression before the start of experiments. Flies were flipped onto fresh medium every third day. Lines obtained from the University of Indiana Bloomington *Drosophila* Stock Center (BDSC) include the following: *Akh-GAL4* (#25684); *AstC::2A::GAL4* (#84595) and *AstC-R1::2A::GAL4* (#84596)^35^; CaLexA system (#66542): *LexAop-CD8::GFP::2A::CD8::GFP; UAS-LexA::VP16::NFAT, LexAop-CD2::GFP/TM6B, Tb*)^28^; *Tub-GAL80*^*TS*^ (#7108); *UAS-Akh* (#27343); *UAS-Cas9*.*P2* (#58985)^56^; *UAS-Tsc1, UAS-Tsc2*^36^ (#80576); and UAS-RNAi lines^57,58^ targeting *AstC-R1* (#27506) and *AstC-R2* (“*AstC-R2-RNAi* ^#2^” #25940). *Tub-GAL80*^*TS*^ and *UAS-Cas9*.*P2* transgenes were recombined onto the same chromosome in-house and combined with *GAL4* lines. The Vienna *Drosophila* Resource Center (VDRC) provided the line *w*^*1118*^ (#60000, isogenic with the VDRC RNAi lines), which crossed to the *GAL4* drivers lines was used as a controls in all experiments, as well as numerous UAS-RNAi lines^59^ listed in Table S1, including lines targeting *AKH* (#105063), *AstC* (#102735), and *AstC-R2* (“*AstC-R2-RNAi*^#1^” #106146). *Ilp2-GAL4, UAS-GFP* (original GAL4 construct from BDSC #37516^60^) was a gift of Pierre Léopold (Institut Pasteur, France). *voilà-GAL4*^61^ was a gift of Alessandro Scopelliti (University of Glasgow, UK). *AstC-R1::2A::GAL4*^23^ and *AstC-R2::2A::GAL4*^23^ were gifts of Shu Kondo (Tohoku University).

### Generation of *UAS-gRNA* lines for tissue-specific CRISPR knockouts

GAL4-inducible *UAS-2xgRNA* constructs allowing tissue-specific disruption of the *AstC* or *AstC-R2* loci were generated in the pCFD6 backbone^62^, obtained from AddGene (#73915; https://addgene.org/). The target sequences (Table S2) were identified using the E-CRISP service^63^ (https://e-crisp.org/E-CRISP/). The *AstC* construct should lead to deletion of most of the peptide-encoding sequence; the *AstC-R2* construct should remove a large part of the locus encoding the transmembrane domains of the receptor, even if the gene retains any function. Putative clones were sequenced, and correct constructs were integrated into the fly genome at the *attP2* (third chromosome) site by BestGene, Inc. (Chino Hills, CA).

### Starvation-survival assays

To assay survival under starvation, flies were transferred without anesthesia to vials of starvation medium (1% agar in water) and kept at 29 °C; the number of dead animals was assessed every 4-8 hours until the completion of the assay. For each genotype, 150 flies were used, except for the rescue experiment with *AKH* overexpression where 60 flies were used. Statistical significance was computed using the Kaplan-Meier log-rank survival functions of the Prism program (GraphPad).

### Immunohistochemistry and confocal imaging

Adult guts and brains were dissected in cold PBS and fixed in 4% paraformaldehyde in PBS at room temperature for 45 or 60 minutes, respectively, with gentle agitation. For corpora cardiaca (CC) preparations, the CCs were slightly pulled out from the head and pre-fixed with head for 30 minutes to minimize variations in the starvation period between different treatments. After pre-fixation, CCs were finely dissected and fixed again with 4% paraformaldehyde in PBS for an additional 10 minutes. Tissues were rinsed with PBST (PBS + 0.1% Triton X-100, Merck #12298) three times for fifteen minutes each, blocked with 3% normal goat serum (Sigma) in PBST at room temperature for one hour, and incubated overnight at 4 °C with gentle agitation in primary antibodies (below) diluted in blocking solution. Primary-antibody solution was removed, and tissues were washed three times with PBST and incubated with secondary antibodies diluted in PBST overnight at 4 °C with gentle agitation, avoiding light. Tissues were washed three times with PBST and mounted in VECTASHIELD mounting medium containing DAPI as counterstain (Vector Laboratories, #H-1200) on slides treated with poly-L-lysine (Sigma, #P8920). Tissues were imaged on a Zeiss LSM-900 confocal microscope using a 20x objective and a 1-micron Z-step in the Zen software package. Post-imaging analysis was performed using the open-source *FIJI* software package^64^. For quantification of AKH and AstC, stacks were Z-projected, AstC-positive cells were identified manually, and staining intensity was measured using the “Measure” tool in FIJI; raw integrated density values per cell were averaged. Antibodies used included rabbit antibody against the processed AKH peptide^24^ (kind gift of Jae Park, U. Tennessee), 1:500; mouse anti-Prospero clone MR1A^65^ (University of Iowa Developmental Studies Hybridoma Bank), 1:20; rabbit anti-DILP2^66^ (kind gift of Ernst Hafen, ETH Zurich), 1:1000; mouse anti-DILP3^14^ (kind gift of Jan Veenstra, University of Bordeaux), 1:500; rabbit anti-AstC^14^ (also kindly given by Jan Veenstra), 1:500; mouse anti-GFP (ThermoFisher #A11120), 1:500; Alexa Fluor 488-conjugated goat anti-mouse (ThermoFisher #A32723), 1:500; Alexa Fluor 488-conjugated goat anti-rabbit (ThermoFisher #A11008), 1:500; and Alexa Fluor 555-conjugated goat anti-mouse (ThermoFisher #A32732), 1:500.

### Binding assay with labeled AstC

BODIPY-labeled AstC was synthesized by Cambridge Peptides, Ltd (UK). The endogenous AstC peptide has the sequence “QVRYRQCYFNPISCF”, cyclized via a disulfide bridge between its cysteines. It is not C-terminally amidated, but the N-terminal residue is processed into a pyroglutamate. Since the C-terminal end of the peptide is more conserved between species and thus likely most important for binding, we sought to label the N terminus. Pyroglutamate lacks a free N-terminal amino group, so it was replaced by a glycine spacer. Thus, the peptide “GGVRYRQCYFNPISCF” was synthesized and cyclized via its cysteine residues; BODIPY-TMR (BODIPY-543) dye was conjugated to the N terminus by an NHS ester linkage. Synthesized peptide was redissolved at 50 μM in 50%/50% DMSO/water. CC complexes were dissected from adult females expressing GFP in the CC (*AKH>GFP* crossed to either *w*^*1118*^ or *UAS-AstC-R2-RNAi #106146*) in artificial hemolymph-like solution^67^ containing 60 mM sucrose (inert osmolyte) and 60 mM trehalose, which mimics a “fed”-like state. Tissues were mounted in artificial hemolymph on poly-lysine-coated glass-bottomed dishes (MatTek, #P35G-0-10-C) and used in a ligand-receptor binding assay as previously described^25^. In brief, labeled peptide was diluted to 1 μM in artificial hemolymph and added at a 1:10 ratio to the mounted tissues (final concentration 100 nM), and bound peptide was imaged using an LSM-900 confocal microscope (Zeiss) in AiryScan2 super-resolution mode with a 40x objective and GFP and BODIPY-543 filter settings.

### Transcript measurement using qPCR

RNA isolation was performed using the RNeasy Mini Plus kit (Qiagen #74136). Tissues (five heads, five guts, or five whole animals per sample, times 6-8 replicate samples per condition or genotype) were homogenized in 2-mL tubes containing lysis buffer plus 1% 2-mercaptoethanol using a TissueLyser LT bead mill (Qiagen) and 5-mm stainless-steel beads (Qiagen #69989) following the manufacturer’s protocol. RNA was reverse-transcribed using the High-Capacity cDNA Synthesis kit (Applied Biosystems, #4368814). QPCR was performed using RealQ Plus 2x Master Mix Green (Ampliqon, #A324402) on Stratagene Mx3005P (Agilent Technologies) and Quant Studio 5 (Applied Biosystems) machines. Results were normalized against expression of *Rp49*, and differences in transcript abundance were computed using the delta-delta-Ct method. The oligos used are listed in Table S2.

### Western blotting

Sets of three adults were homogenized in 100 μl 2x SDS sample buffer (Bio-Rad #1610737) containing 5% (355 mM) 2-mercaptoethanol, heated at 95° for 5 minutes, and centrifuged at maximum speed for 5 minutes to pellet debris. Fifteen microliters of each supernatant was loaded into a precast 4%-20% polyacrylamide gradient gel (Bio-Rad). After electrophoresis, proteins were transferred to polyvinylidene difluoride membrane (PVDF, Millipore) using the Trans-Blot Turbo Transfer Pack (Bio-Rad #1704158) on the Trans-Blot Turbo apparatus (Bio-Rad). The membrane was blocked in blocking buffer (LI-COR, #927-40100) and incubated with rabbit anti-phospho-Akt (Cell Signaling Technology, #4054) and mouse anti-histone H3 (Abcam, #ab1791), both diluted 1:1000 in blocking buffer + 0.2% Tween-20. The membrane was rinsed and incubated with IRDye 680RD anti-mouse (LI-COR #925-68070) and IRDye 800CW anti-rabbit (LI-COR #925-32210), diluted 1:10,000; bands were visualized using the Odyssey Fc imaging system (LI-COR). The membrane was stripped, and total Akt was stained using a similar procedure with rabbit anti-Akt (Cell Signaling Technology, #4691, 1:1000).

### Feeding assays

Short-term food intake was measured using a dye-feeding assay^68,69^. At the time of the main morning meal (after lights-on in the incubator), flies were quickly transferred to nutrient-balanced sugar-yeast food^70^ (90 g/L sucrose, 80 g/L yeast, 10 g/L agar, supplemented with 1 mL/L propionic acid and 1 g/L methyl-4-hydroxybenzoate) containing 0.5% erioglaucine dye (Sigma-Aldrich, #861146) and allowed to feed for 30 minutes. Other flies were allowed to feed on undyed medium for use in baselining spectrophotometric measurements. Ten sets of 3 animals for each genotype were homogenized in 150 μl PBS, pH 7.5, using a TissueLyser LT (Qiagen) bead mill with 5-mm stainless-steel beads (Qiagen, #69989). Insoluble debris was pelleted by centrifugation, 50 μl of the supernatant was transferred to a 384-well plate well, and absorbance at 629 nm was measured using an Ensight multi-mode plate reader (PerkinElmer). Readings were calibrated against an erioglaucine standard curve to compute the amount of food consumed per fly for 30 min. Longer-term food consumption was monitored using the CAFE assay^30^. Each fly was placed into a 2-ml Eppendorf tube with a 5-μl microcapillary inserted through a hole in the lid; the capillary was filled with liquid sugar-yeast medium^30^ (50 g/L sucrose, 50 g/L yeast extract, supplemented with 1 mL/L propionic acid and 1 g/L methyl-4-hydroxybenzoate). To minimize evaporation, capillary-equipped tubes were kept in a moist chamber, and fly-free tubes were used to control for the level of evaporation. The amount of food consumed was determined by measuring the food level within the capillary tube every 24 hours for 3 days. Fifteen flies were individually scored for each genotype. To monitor the performance of feeding-related behaviors, rather than the consumed amounts, we used the FLIC assay^31^. *Drosophila* Feeding Monitors (Sable Systems, US) were kept in a 29-degree incubator at 70% humidity with a 12-hour light cycle. Individual flies were placed in the monitors the night before the assay to allow them to acclimate; the following subjective dawn, fresh sugar-water medium (5% sucrose) was added, and feeding behaviors were recorded for 24 hours. Physical contact between the flies and the sugar-water feeding medium were recorded using the manufacturer’s software and analyzed by a package in the *R* analysis environment (https://github.com/PletcherLab/FLIC_R_Code)^31^.

### Sleep assays

Sleep assays were carried out using the *Drosophila* Activity Monitor system (TriKinetics, Waltham, MA). Individual male or female flies (8 days old: 3 days at 18 degrees post-eclosion to complete the transition to adult physiology, followed by five days of 29-degree pretreatment to induce RNAi) were transferred to 2-mm inner-diameter glass tubes mounted in DAM2 monitors; one end of each tube was sealed using a 250-μL individual PCR tube filled with sugar food (5% sucrose and 1% agar in water), and the other end was plugged with plastic foam. Monitors were placed within a 29-degree incubator (60% humidity) with a 12-hour light/dark cycle. The first day of data, reflecting behavioral acclimation to the environment, was discarded. Activity was recorded for 24 hours from subjective dawn; at the second dawn, the food-containing PCR tubes were replaced with similar tubes containing only 1% agar, and the subsequent 24 hours of behavior under starvation was recorded. “Sleep” was defined as five minutes of quiescence. Data were analyzed using pySolo^71^ and custom scripts written in the MATLAB environment (The MathWorks, Natick, MA).

### Metabolite measurements

Bulk tri-/-diacylglycerides and glycogen were measured using protocols described in detail previously^70,72^. For each sample, 4 adult flies were homogenized in PBS buffer with 0.5% Tween-20 (Sigma, #1379) in a TissueLyser LT (Qiagen) bead mill with 5-mm stainless-steel beads. An aliquot of homogenate was used for bicinchonic-acid protein-level determination using commercial components (Sigma, #B9643, #C2284, and #P0914). Glycogen content was assayed by hydrolyzing glycogen into glucose with amyloglucosidase (Sigma, #A7420) and measuring the liberated glucose using a colorimetric assay kit (Sigma, #GAGO20). For determination of circulating glucose, hemolymph was extracted^73^ from adult females before and after starvation, and glucose was measured using the colorimetric assay (Sigma, #GAGO20). Tri- and diacylglycerides were measured by cleaving their ester bonds using triglyceride reagent (Sigma, #T2449) to liberate glycerol, which was measured using Free Glycerol Reagent (Sigma, #F6428) in a colorimetric assay. Absorbances from both assays were read at 540 nm using an Ensight multimode plate reader (PerkinElmer). Concentrations of TAG and glycogen were quantified using corresponding glucose and glycerol standard curves.

### Statistics

All statistics were computed using Prism software (GraphPad). Starvation-survival curves were analyzed using Kaplan-Meier log-rank tests. Multiple comparisons were analyzed using ANOVA and Kruskal-Wallis test, and pairwise comparisons were made using t-test for normally distributed data or nonparametric Mann-Whitney U test. Significance is shown in figures as *: P<0.05; **: P<0.01; ***: P<0.001.

## Acknowledgements

Anti-Akh was a generous gift of Jae Park (University of Tennessee). Anti-DILP2 was a kind gift of Ernst Hafen (ETH Zurich). Anti-AstC and anti-DILP3 were kind gifts of Jan Veenstra (University of Bordeaux). We thank the University of Indiana Bloomington *Drosophila* Stock Center and the Vienna Drosophila *Resource* Center for maintaining and providing fly lines, the University of Iowa Developmental Studies Hybridoma bank for producing and providing anti-Prospero, and AddGene for producing and providing pCFD6. This work was supported by Novo Nordisk Foundation grant 0054632 and Lundbeck Foundation grant 2019-772 to K.R. T.K., M.T.N., and K.A.H. were supported by funding from the Villum Foundation (15365) and Danish Council for Independent Research Natural Sciences (9064-00009B) to K.A.H.

## Author contributions

M.J.T., S.N., and K.R. conceived and designed the study. O.K., L.N., N.A., T.K., A.M., S.N., K.A.H., M.J.T., and K.R. designed, performed, and analyzed experiments. M.T.N. performed experiments. M.L. analyzed data. M.J.T. and K.R. wrote the manuscript.

## Supplemental Material

**Figure S1.**
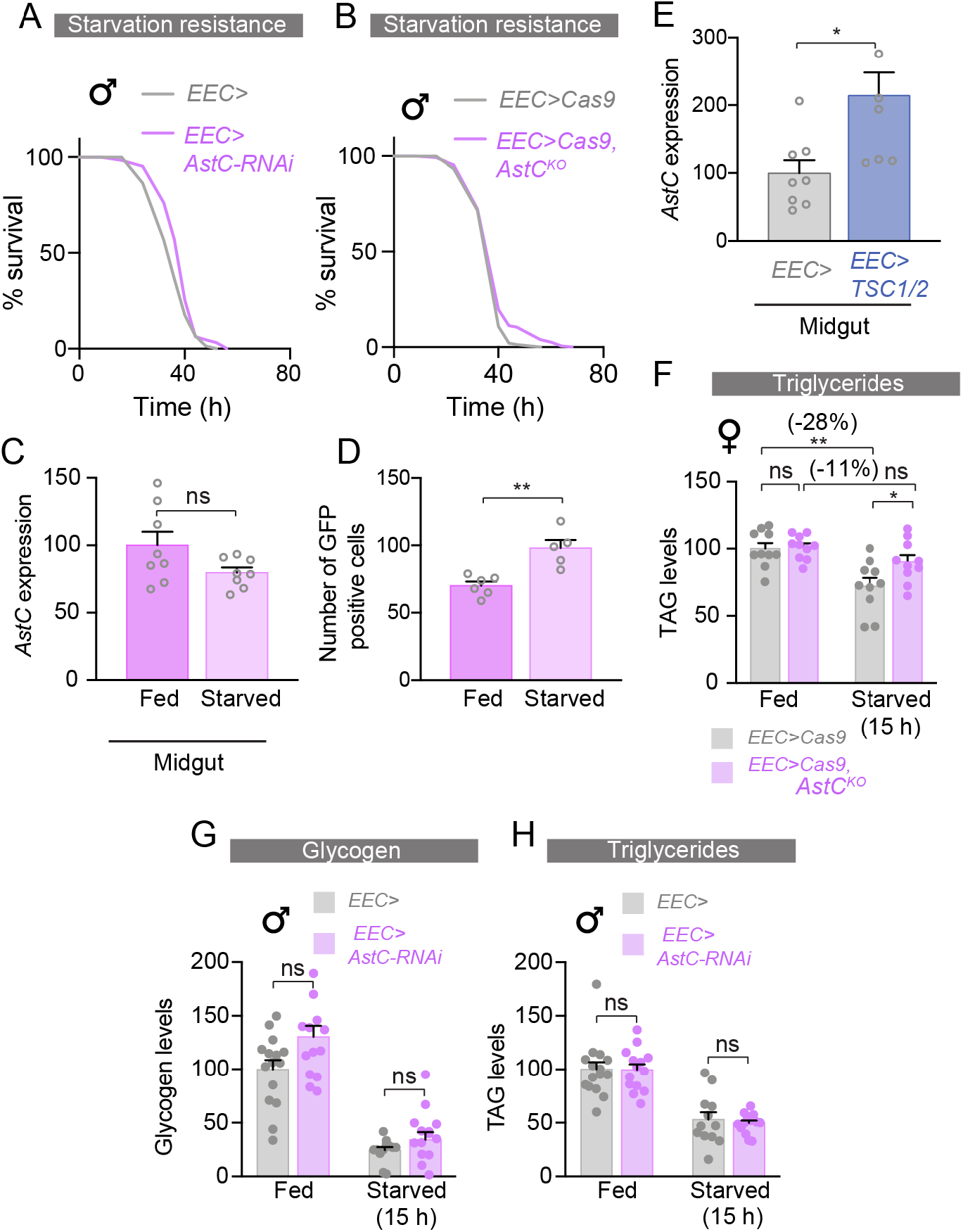
Gut-derived AstC does not alter starvation survival or starvation-induced energy mobilization in males; *AstC* is regulated by TOR signaling in females. **(A and B)** Male flies with *AstC*-targeting RNAi **(A)** or CRISPR knockout **(B)** in their EECs (magenta) do not exhibit longer survival during starvation than controls (gray). **(C)** *AstC* transcript levels in the female gut are not altered by 6-hour starvation. **(D)** The number of *AstC-GAL4*-expressing cells exhibiting observable CaLexA>GFP in the female gut is increased by starvation (results represent analysis of guts from 6 individual animals). (**E)** Inhibition of TOR via expressing *TSC1* and *TSC2* in adult female EECs for six hours (via temperature induction) increases *AstC* transcript levels in dissected female guts. **(F)** CRISPR-mediated knockout of *AstC* in the EECs recapitulates the lipid-mobilization defect of *AstC-RNAi* in females. **(G and H**) *AstC* RNAi in the EECs does not alter the mobilization of glycogen **(G)** or TAGs **(H)** in males. Error bars indicate standard error of the mean (SEM). ns, non-significant; *, P<0.05; and **, P<0.01.

**Figure S2.**
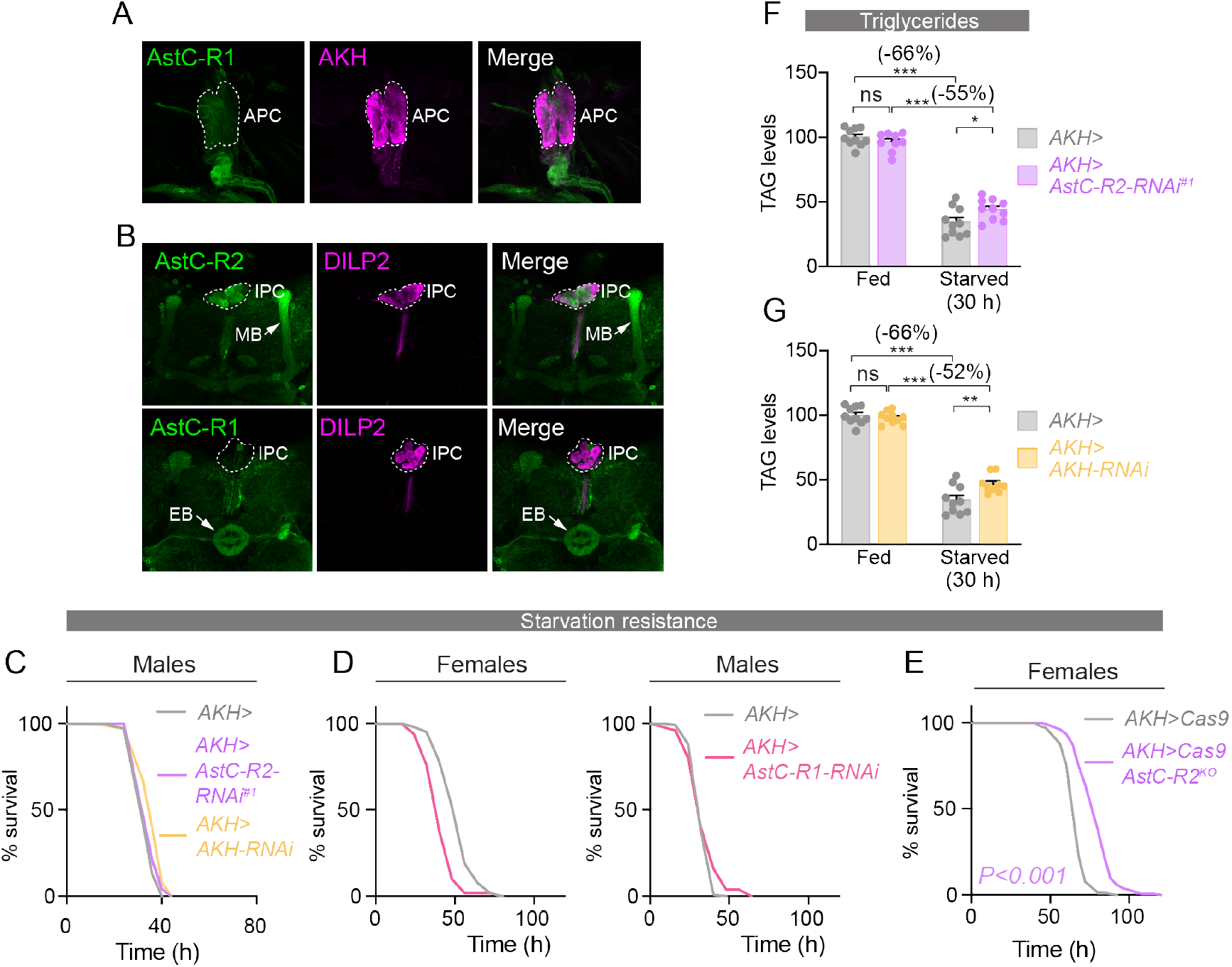
AstC effects are mediated by AstC-R2 in the APCs, not by effects on the IPCs. **(A)** *AstC-R1::2A::GAL4, UAS-GFP* (green) does not strongly mark the female CC (marked by anti-AKH, magenta). **(B)** *AstC-R2>GFP* (green) reports AstC-R2 expression in the IPCs (marked by anti-DILP2, magenta) and the mushroom bodies (“MB”); *AstC-R1>GFP* reports AstC-R1 expression in the ellipsoid body (“EB”) of the central complex of the brain, but not in the IPCs (anti-DILP2, magenta). **(C)** Knockdown of *AstC-R2* in the APCs (magenta) does not prolong male survival relative to controls (gray). Knockdown of *AKH* itself in the male APCs also has no effect on starvation survival (orange). **(D)** Knockdown of *AstC-R1* in the APCs (rose) does not prolong starvation in survival in females (left panel) or males (right panel), compared with controls (gray). **(E)** CRISPR-mediated deletion of *AstC-R2* (magenta) in the female APCs phenocopies the starvation-survival effect of *AstC-R2-RNAi* in this tissue. **(F and G, above C-E)** RNAi against *AstC-R2* (**F**, magenta) or *Akh* (**G**, orange) in the female APCs has similar suppressive effects on TAG mobilization during 30-hour starvation. Error bars indicate standard error of the mean (SEM). ns, non-significant; *, P<0.05; **, P<0.01; and ***, P<0.001.

**Figure S3.**
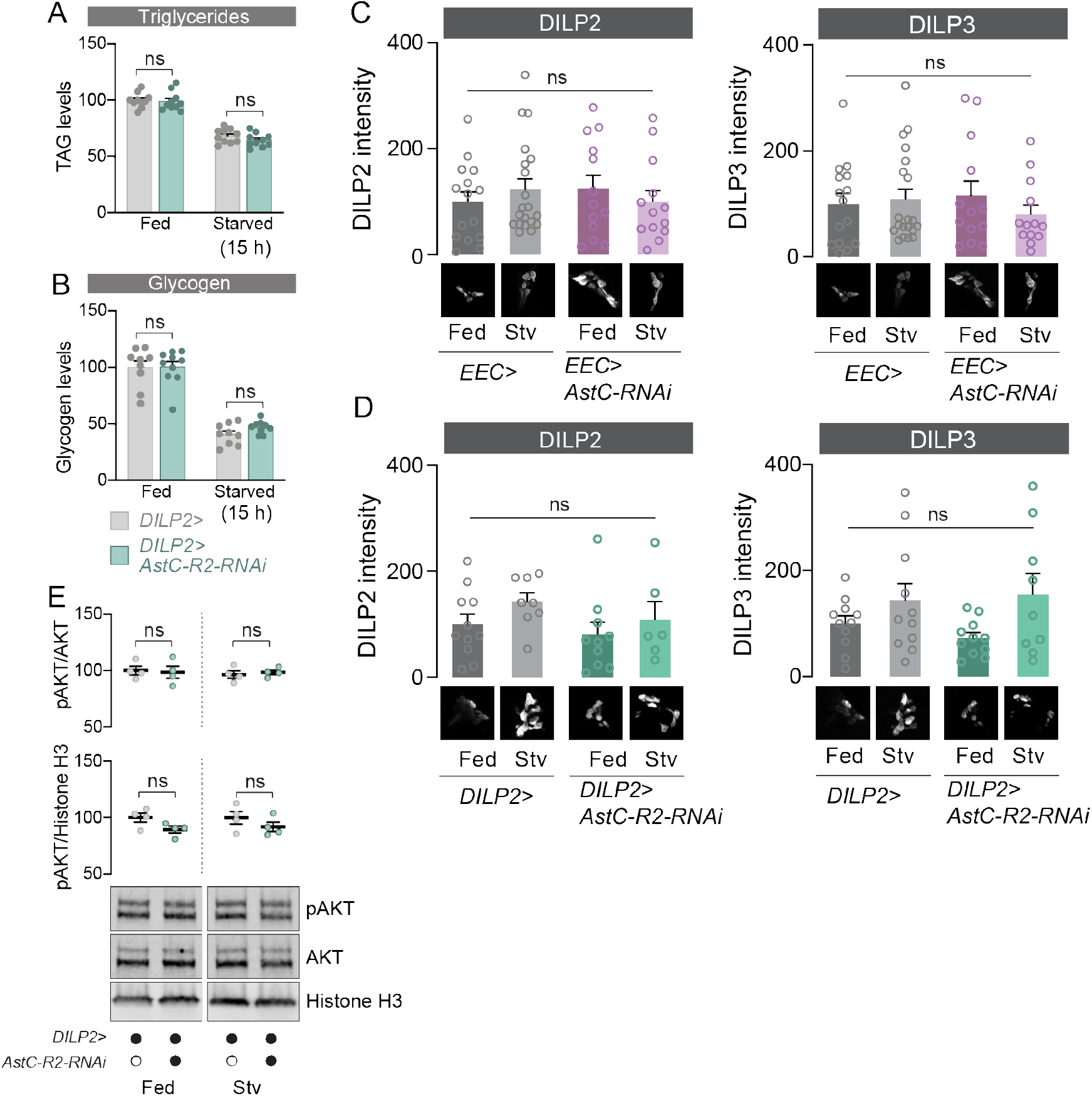
Although AstC-R2 appears to be expressed in the IPCs, it does not affect insulin signaling under the nutrient-stress conditions examined. **(A and B)** Knockdown of *AstC-R2* in the IPCs (teal) does not alter the mobilization of glycogen **(A)** or TAGs **(B)** during 15-hour starvation. **(C)** Knockdown of *AstC* in the EECs does not alter the accumulation of DILP2 (left panel) or DILP3 (right panel) in female IPCs, as measured by anti-DILP immunostaining. **(D)** Likewise, knockdown of *AstC-R2* in the IPCs does not strongly alter the accumulation of DILP2 (left) or DILP3 (right) in these cells. **(E)** Knockdown of *AstC-R2* in the IPCs does not strongly alter peripheral insulin-signaling activity during feeding or starvation, as measured by anti-phospho-AKT Western blotting of whole females. Top: anti-phospho-AKT (pAKT) staining levels normalized against total AKT staining; middle: anti-pAKT staining normalized against histone H3 staining; bottom: Western-blot lanes illustrating similar staining intensities under all conditions. Error bars indicate standard error of the mean (SEM). ns, non-significant.

**Figure S4.**
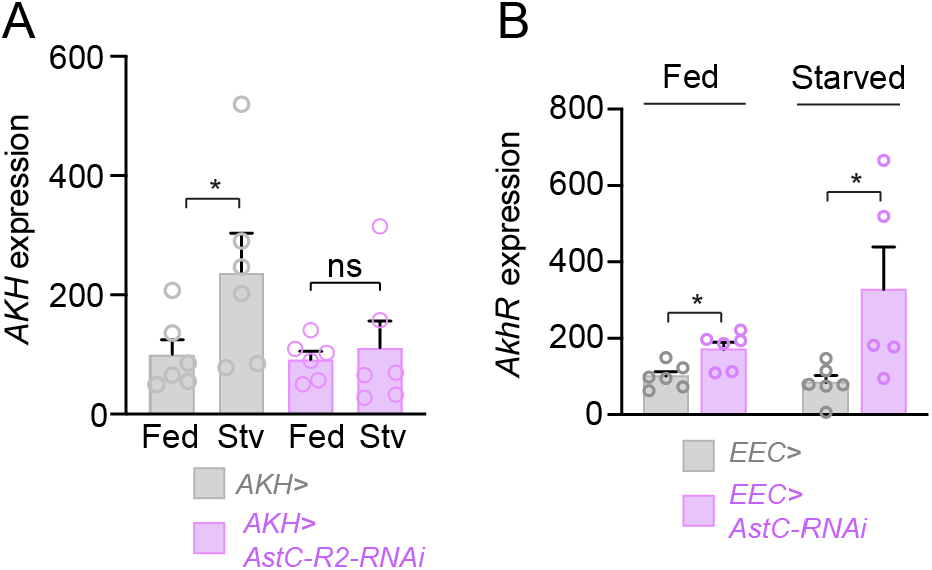
Transcript levels of *AKH* and *AkhR* are altered with disruption of AstC signaling. **(A)** In control animals (gray), *AKH* transcript levels are increased with starvation; this upregulation does not occur, however, when *AstC-R2* is knocked down in the APCs (magenta). **(B)** Expression of *AkhR* is higher in animals in which EEC expression of *AstC* is knocked down (magenta). Error bars indicate standard error of the mean (SEM). *, P<0.05.

**Figure S5.**
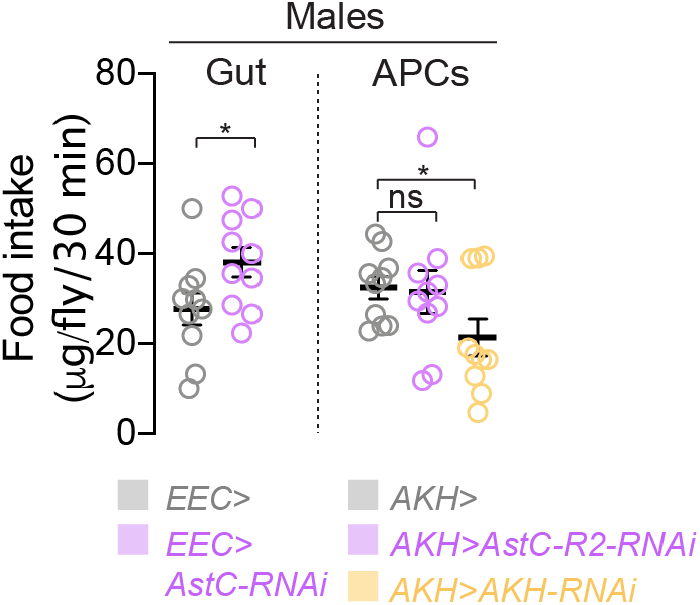
Effects of gut AstC and AKH on feeding in males. Food intake over 30 minutes, measured by a dye-feeding assay in males expressing RNAi against *AstC* in the EECs (magenta) and controls (grey). Error bars indicate standard error of the mean (SEM). ns, non-significant; *, P<0.05.

**Table S1.**
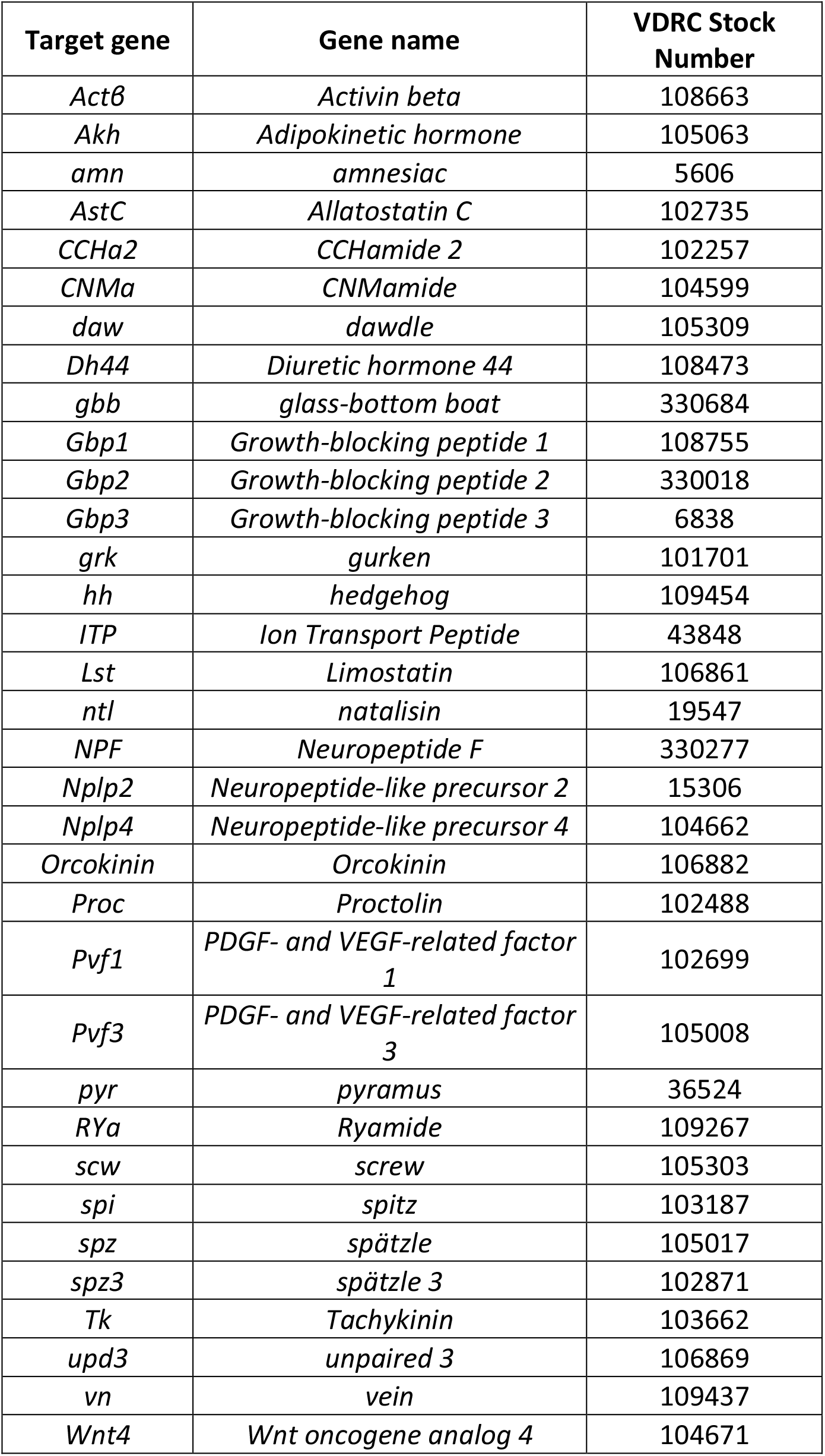
Transgenic RNAi lines used for screening.

**Table S2.**
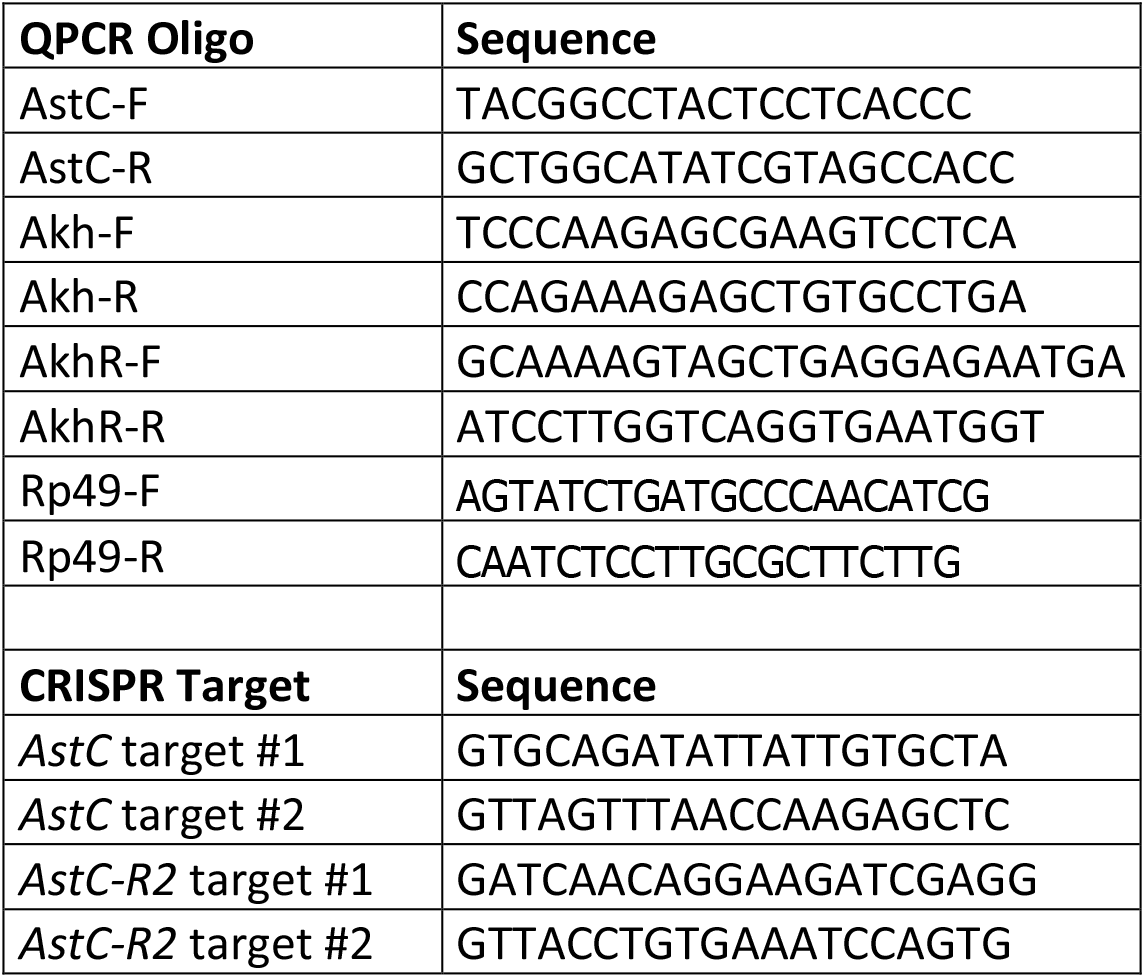
Sequences of qPCR primers and gRNA target sites.

## References

1 Koyama, T., Texada, M. J., Halberg, K. A. & Rewitz, K. Metabolism and growth adaptation to environmental conditions in Drosophila. Cell Mol Life Sci, doi:10.1007/s00018-020-03547-2 (2020).

2 Rehfeld, J. F. The new biology of gastrointestinal hormones. Physiol Rev 78, 1087–1108, doi:10.1152/physrev.1998.78.4.1087 (1998).

3 Gribble, F. M. & Reimann, F. Function and mechanisms of enteroendocrine cells and gut hormones in metabolism. Nat Rev Endocrinol 15, 226–237, doi:10.1038/s41574-019-0168-8 (2019).

4 Inui, A. Ghrelin: an orexigenic and somatotrophic signal from the stomach. Nat Rev Neurosci 2, 551–560, doi:10.1038/35086018 (2001).

5 Nauck, M. A. & Meier, J. J. Incretin hormones: Their role in health and disease. Diabetes Obes Metab 20 Suppl 1, 5–21, doi:10.1111/dom.13129 (2018).

6 Colombani, J. & Andersen, D. S. The Drosophila gut: A gatekeeper and coordinator of organism fitness and physiology. Wiley Interdiscip Rev Dev Biol, e378, doi:10.1002/wdev.378 (2020).

7 Miguel-Aliaga, I., Jasper, H. & Lemaitre, B. Anatomy and Physiology of the Digestive Tract of Drosophila melanogaster. Genetics 210, 357–396, doi:10.1534/genetics.118.300224 (2018).

8 Texada, M. J., Koyama, T. & Rewitz, K. Regulation of Body Size and Growth Control. Genetics 216, 269–313, doi:10.1534/genetics.120.303095 (2020).

9 Wang, S., Tulina, N., Carlin, D. L. & Rulifson, E. J. The origin of islet-like cells in Drosophila identifies parallels to the vertebrate endocrine axis. Proc Natl Acad Sci U S A 104, 19873–19878, doi:10.1073/pnas.0707465104 (2007).

10 Bharucha, K. N., Tarr, P. & Zipursky, S. L. A glucagon-like endocrine pathway in Drosophila modulates both lipid and carbohydrate homeostasis. J Exp Biol 211, 3103–3110, doi:10.1242/jeb.016451 (2008).

11 Yu, Y. et al. Regulation of starvation-induced hyperactivity by insulin and glucagon signaling in adult Drosophila. Elife 5, doi:10.7554/eLife.15693 (2016).

12 Hung, R. J. et al. A cell atlas of the adult Drosophila midgut. Proc Natl Acad Sci U S A 117, 1514–1523, doi:10.1073/pnas.1916820117 (2020).

13 Reiher, W. et al. Peptidomics and peptide hormone processing in the Drosophila midgut. J Proteome Res 10, 1881–1892, doi:10.1021/pr101116g (2011).

14 Veenstra, J. A., Agricola, H. J. & Sellami, A. Regulatory peptides in fruit fly midgut. Cell Tissue Res 334, 499–516, doi:10.1007/s00441-008-0708-3 (2008).

15 Veenstra, J. A. Peptidergic paracrine and endocrine cells in the midgut of the fruit fly maggot. Cell Tissue Res 336, 309–323, doi:10.1007/s00441-009-0769-y (2009).

16 Song, W., Veenstra, J. A. & Perrimon, N. Control of lipid metabolism by tachykinin in Drosophila. Cell Rep 9, 40–47, doi:10.1016/j.celrep.2014.08.060 (2014).

17 Ameku, T. et al. Midgut-derived neuropeptide F controls germline stem cell proliferation in a mating-dependent manner. PLoS Biol 16, e2005004, doi:10.1371/journal.pbio.2005004 (2018).

18 Scopelliti, A. et al. A Neuronal Relay Mediates a Nutrient Responsive Gut/Fat Body Axis Regulating Energy Homeostasis in Adult Drosophila. Cell Metab, doi:10.1016/j.cmet.2018.09.021 (2018).

19 Benguettat, O. et al. The DH31/CGRP enteroendocrine peptide triggers intestinal contractions favoring the elimination of opportunistic bacteria. PLoS Pathog 14, e1007279, doi:10.1371/journal.ppat.1007279 (2018).

20 Veenstra, J. A. Allatostatin C and its paralog allatostatin double C: the arthropod somatostatins. Insect Biochem Mol Biol 39, 161–170, doi:10.1016/j.ibmb.2008.10.014 (2009).

21 Wat, L. W. et al. A role for triglyceride lipase brummer in the regulation of sex differences in Drosophila fat storage and breakdown. PLoS Biol 18, e3000595, doi:10.1371/journal.pbio.3000595 (2020).

22 Diaz, M. M., Schlichting, M., Abruzzi, K. C., Long, X. & Rosbash, M. Allatostatin-C/AstC-R2 Is a Novel Pathway to Modulate the Circadian Activity Pattern in Drosophila. Curr Biol 29, 13–22 e13, doi:10.1016/j.cub.2018.11.005 (2019).

23 Kondo, S. et al. Neurochemical Organization of the Drosophila Brain Visualized by Endogenously Tagged Neurotransmitter Receptors. Cell Rep 30, 284–297 e285, doi:10.1016/j.celrep.2019.12.018 (2020).

24 Lee, G. & Park, J. H. Hemolymph sugar homeostasis and starvation-induced hyperactivity affected by genetic manipulations of the adipokinetic hormone-encoding gene in Drosophila melanogaster. Genetics 167, 311–323 (2004).

25 Halberg, K. A., Terhzaz, S., Cabrero, P., Davies, S. A. & Dow, J. A. Tracing the evolutionary origins of insect renal function. Nature communications 6, 6800, doi:10.1038/ncomms7800 (2015).

26 Quesada, I., Tuduri, E., Ripoll, C. & Nadal, A. Physiology of the pancreatic alpha-cell and glucagon secretion: role in glucose homeostasis and diabetes. J Endocrinol 199, 5–19, doi:10.1677/JOE-08-0290 (2008).

27 Kim, S. K. & Rulifson, E. J. Conserved mechanisms of glucose sensing and regulation by Drosophila corpora cardiaca cells. Nature 431, 316–320, doi:10.1038/nature02897 nature02897 [pii] (2004).

28 Masuyama, K., Zhang, Y., Rao, Y. & Wang, J. W. Mapping neural circuits with activity-dependent nuclear import of a transcription factor. Journal of neurogenetics 26, 89–102, doi:10.3109/01677063.2011.642910 (2012).

29 Hirsch, I. B. et al. Insulin and glucagon in prevention of hypoglycemia during exercise in humans. Am J Physiol 260, E695–704, doi:10.1152/ajpendo.1991.260.5.E695 (1991).

30 Ja, W. W. et al. Prandiology of Drosophila and the CAFE assay. Proc Natl Acad Sci U S A 104, 8253–8256, doi:10.1073/pnas.0702726104 (2007).

31 Ro, J., Harvanek, Z. M. & Pletcher, S. D. FLIC: high-throughput, continuous analysis of feeding behaviors in Drosophila. PLoS One 9, e101107, doi:10.1371/journal.pone.0101107 (2014).

32 Galikova, M., Klepsatel, P., Xu, A. & Kuhnlein, R. P. The obesity-related Adipokinetic hormone controls feeding and expression of neuropeptide regulators of Drosophila metabolism. Eur J Lipid Sci Technol 119, 1600138 (2017).

33 Jourjine, N., Mullaney, B. C., Mann, K. & Scott, K. Coupled Sensing of Hunger and Thirst Signals Balances Sugar and Water Consumption. Cell 166, 855–866, doi:10.1016/j.cell.2016.06.046 (2016).

34 Park, J. H. et al. A subset of enteroendocrine cells is activated by amino acids in the Drosophila midgut. FEBS Lett 590, 493–500, doi:10.1002/1873-3468.12073 (2016).

35 Deng, B. et al. Chemoconnectomics: Mapping Chemical Transmission in Drosophila. Neuron 101, 876–893 e874, doi:10.1016/j.neuron.2019.01.045 (2019).

36 Tapon, N., Ito, N., Dickson, B. J., Treisman, J. E. & Hariharan, I. K. The Drosophila tuberous sclerosis complex gene homologs restrict cell growth and cell proliferation. Cell 105, 345–355 (2001).

37 Chen, J., Kim, S. M. & Kwon, J. Y. A Systematic Analysis of Drosophila Regulatory Peptide Expression in Enteroendocrine Cells. Molecules and cells 39, 358–366, doi:10.14348/molcells.2016.0014 (2016).

38 Dutta, D., Buchon, N., Xiang, J. & Edgar, B. A. Regional Cell Specific RNA Expression Profiling of FACS Isolated Drosophila Intestinal Cell Populations. Curr Protoc Stem Cell Biol 34, 2F 2 1–2F 2 14, doi:10.1002/9780470151808.sc02f02s34 (2015).

39 Borbely, A. A. Sleep in the rat during food deprivation and subsequent restitution of food. Brain Res 124, 457–471, doi:10.1016/0006-8993(77)90947-7 (1977).

40 Gylfe, E. & Gilon, P. Glucose regulation of glucagon secretion. Diabetes Res Clin Pract 103, 1–10, doi:10.1016/j.diabres.2013.11.019 (2014).

41 Unger, R. H. & Cherrington, A. D. Glucagonocentric restructuring of diabetes: a pathophysiologic and therapeutic makeover. J Clin Invest 122, 4–12, doi:10.1172/JCI60016 (2012).

42 Rutter, G. A. Regulating glucagon secretion: somatostatin in the spotlight. Diabetes 58, 299–301, doi:10.2337/db08-1534 (2009).

43 Mani, B. K. & Zigman, J. M. A Strong Stomach for Somatostatin. Endocrinology 156, 3876–3879, doi:10.1210/en.2015-1756 (2015).

44 Francis, B. H., Baskin, D. G., Saunders, D. R. & Ensinck, J. W. Distribution of somatostatin-14 and somatostatin-28 gastrointestinal-pancreatic cells of rats and humans. Gastroenterology 99, 1283–1291, doi:10.1016/0016-5085(90)91151-u (1990).

45 Ludington, W. B. & Ja, W. W. Drosophila as a model for the gut microbiome. PLoS Pathog 16, e1008398, doi:10.1371/journal.ppat.1008398 (2020).

46 Worthington, J. J., Reimann, F. & Gribble, F. M. Enteroendocrine cells-sensory sentinels of the intestinal environment and orchestrators of mucosal immunity. Mucosal Immunol 11, 3–20, doi:10.1038/mi.2017.73 (2018).

47 Bachtel, N. D., Hovsepian, G. A., Nixon, D. F. & Eleftherianos, I. Allatostatin C modulates nociception and immunity in Drosophila. Sci Rep 8, 7501, doi:10.1038/s41598-018-25855-1 (2018).

48 An, S. et al. Insect neuropeptide bursicon homodimers induce innate immune and stress genes during molting by activating the NF-kappaB transcription factor Relish. PLoS One 7, e34510, doi:10.1371/journal.pone.0034510 (2012).

49 Clegg, D. J. & Mauvais-Jarvis, F. An integrated view of sex differences in metabolic physiology and disease. Mol Metab 15, 1–2, doi:10.1016/j.molmet.2018.06.011 (2018).

50 Karastergiou, K., Smith, S. R., Greenberg, A. S. & Fried, S. K. Sex differences in human adipose tissues - the biology of pear shape. Biol Sex Differ 3, 13, doi:10.1186/2042-6410-3-13 (2012).

51 Jensen, K., McClure, C., Priest, N. K. & Hunt, J. Sex-specific effects of protein and carbohydrate intake on reproduction but not lifespan in Drosophila melanogaster. Aging Cell 14, 605–615, doi:10.1111/acel.12333 (2015).

52 Efeyan, A., Comb, W. C. & Sabatini, D. M. Nutrient-sensing mechanisms and pathways. Nature 517, 302–310, doi:10.1038/nature14190 (2015).

53 Gonzalez, A. & Hall, M. N. Nutrient sensing and TOR signaling in yeast and mammals. EMBO J 36, 397–408, doi:10.15252/embj.201696010 (2017).

54 Baggio, L. L. & Drucker, D. J. Biology of incretins: GLP-1 and GIP. Gastroenterology 132, 2131–2157, doi:10.1053/j.gastro.2007.03.054 (2007).

55 Breen, D. M., Rasmussen, B. A., Cote, C. D., Jackson, V. M. & Lam, T. K. Nutrient-sensing mechanisms in the gut as therapeutic targets for diabetes. Diabetes 62, 3005–3013, doi:10.2337/db13-0523 (2013).

56 Port, F., Chen, H. M., Lee, T. & Bullock, S. L. Optimized CRISPR/Cas tools for efficient germline and somatic genome engineering in Drosophila. Proc Natl Acad Sci U S A 111, E2967–2976, doi:10.1073/pnas.1405500111 (2014).

57 Ni, J. Q. et al. A genome-scale shRNA resource for transgenic RNAi in Drosophila. Nature methods 8, 405–407, doi:10.1038/nmeth.1592 (2011).

58 Perkins, L. A. et al. The Transgenic RNAi Project at Harvard Medical School: Resources and Validation. Genetics 201, 843–852, doi:10.1534/genetics.115.180208 (2015).

59 Dietzl, G. et al. A genome-wide transgenic RNAi library for conditional gene inactivation in Drosophila. Nature 448, 151–156, doi:nature05954 [pii] 10.1038/nature05954 (2007).

60 Rulifson, E. J., Kim, S. K. & Nusse, R. Ablation of insulin-producing neurons in flies: growth and diabetic phenotypes. Science 296, 1118–1120, doi:10.1126/science.1070058296/5570/1118 [pii] (2002).

61 Balakireva, M., Gendre, N., Stocker, R. F. & Ferveur, J. F. The genetic variant Voila causes gustatory defects during Drosophila development. J Neurosci 20, 3425–3433 (2000).

62 Port, F. & Bullock, S. L. Augmenting CRISPR applications in Drosophila with tRNA-flanked sgRNAs. Nat Methods 13, 852–854, doi:10.1038/nmeth.3972 (2016).

63 Heigwer, F., Kerr, G. & Boutros, M. E-CRISP: fast CRISPR target site identification. Nat Methods 11, 122–123, doi:10.1038/nmeth.2812 (2014).

64 Schindelin, J. et al. Fiji: an open-source platform for biological-image analysis. Nature methods 9, 676–682, doi:10.1038/nmeth.2019 (2012).

65 Campbell, G. et al. RK2, a glial-specific homeodomain protein required for embryonic nerve cord condensation and viability in Drosophila. Development 120, 2957–2966 (1994).

66 Bader, R. et al. The IGFBP7 homolog Imp-L2 promotes insulin signaling in distinct neurons of the Drosophila brain. J Cell Sci 126, 2571–2576, doi:10.1242/jcs.120261 (2013).

67 Feng, Y., Ueda, A. & Wu, C. F. A modified minimal hemolymph-like solution, HL3.1, for physiological recordings at the neuromuscular junctions of normal and mutant Drosophila larvae. J Neurogenet 18, 377–402, doi:10.1080/01677060490894522 (2004).

68 Wong, R., Piper, M. D., Wertheim, B. & Partridge, L. Quantification of food intake in Drosophila. PLoS One 4, e6063, doi:10.1371/journal.pone.0006063 (2009).

69 Skorupa, D. A., Dervisefendic, A., Zwiener, J. & Pletcher, S. D. Dietary composition specifies consumption, obesity, and lifespan in Drosophila melanogaster. Aging Cell 7, 478–490, doi:10.1111/j.1474-9726.2008.00400.x (2008).

70 Tennessen, J. M., Barry, W. E., Cox, J. & Thummel, C. S. Methods for studying metabolism in Drosophila. Methods 68, 105–115, doi:10.1016/j.ymeth.2014.02.034 (2014).

71 Gilestro, G. F. & Cirelli, C. pySolo: a complete suite for sleep analysis in Drosophila. Bioinformatics 25, 1466–1467, doi:10.1093/bioinformatics/btp237 (2009).

72 Hildebrandt, A., Bickmeyer, I. & Kuhnlein, R. P. Reliable Drosophila body fat quantification by a coupled colorimetric assay. PLoS One 6, e23796, doi:10.1371/journal.pone.0023796 (2011).

73 Tennessen, J. M., Barry, W. E., Cox, J. & Thummel, C. S. Methods for studying metabolism in Drosophila. Methods 68, 105–115, doi:10.1016/j.ymeth.2014.02.034 (2014).

